# A Markovian dynamics for *C. elegans* behavior across scales

**DOI:** 10.1101/2023.10.19.563098

**Authors:** Antonio C. Costa, Tosif Ahamed, David Jordan, Greg J. Stephens

## Abstract

How do we capture the breadth of behavior in animal movement, from rapid body twitches to aging? Using high-resolution videos of the nematode worm *C. elegans*, we show that a single dynamics connects posture-scale fluctuations with trajectory diffusion, and longer-lived behavioral states. We take short posture sequences as an instantaneous behavioral measure, fixing the sequence length for maximal prediction. Within the space of posture sequences we construct a fine-scale, maximum entropy partition so that transitions among microstates define a high-fidelity Markov model, which we also use as a means of principled coarse-graining. We translate these dynamics into movement using resistive force theory, capturing the statistical properties of foraging trajectories. Predictive across scales, we leverage the longest-lived eigenvectors of the inferred Markov chain to perform a top-down subdivision of the worm’s foraging behavior, revealing both “runs-and-pirouettes” as well as previously uncharacterized finer-scale behaviors. We use our model to investigate the relevance of these fine-scale behaviors for foraging success, recovering a trade-off between local and global search strategies.

**SIGNIFICANCE STATEMENT:** Complex phenotypes, such as an animal’s behavior, generally depend on an overwhelming number of processes that span a vast range of scales. While there is no reason that behavioral dynamics permit simple models, by subsuming inherent nonlinearities and memory into maximally-predictive microstates, we find one for *C. elegans* foraging. The resulting “Markov worm” is effectively indistinguishable from real worm motion across a range of timescales, and we can decompose our model dynamics both to recover and discover behavioral states. Employing a simple form of substrate interactions, we connect postures to trajectories, illuminating how worms explore the environment. In more complex organisms, our approach can also link behaviors across time, from rapid muscular control to neuromodulation.

## INTRODUCTION

From molecular motors contracting muscles, to neurons processing an ever changing environment, or the large-scale diffusion of hormones and other neuromodulatory chemicals, animal behavior arises from biological activity across innumerable spatial and temporal scales. With an instantaneous snapshot of all of these variables, the future behavioral state of the animal would be uniquely defined, a biological setting for the demon of Laplace (see e.g. [1]). Of course, such an approach is practically unrealizable. We are limited to a much smaller set of observations and the unobserved degrees of freedom will generally induce non-Markovianity, or memory, to the dynamics of the variables that we do measure [2, 3]. In animal behavior, interpretations of this memory guide our understanding of the complexity of the process [4, 5]. But what if we could use our observations to construct memory-full state variables that admit predictive, yet minimal-memory dynamics [6, 7]?

The construction of such dynamics appears daunting. We may even conclude that this is impossible if it were not for the fact that it is done routinely in physical systems. Indeed, it is often the case that a subset of observable functions is enough to capture behavior at a particular scale. Hydrodynamics, for example, can be formulated with effective variables such as fluid velocity, density, or temperature: their memory only coming from the previous state. In behavior, we expect the emergent reconstructed dynamics to be generally high-dimensional in order to account for the multitude of unobserved mechanisms. Yet our approach also suggests a principled coarse-graining. Since the dynamics of the reconstructed states are Markovian, the emergent timescales of the (nonlinear) system are naturally ordered by the eigenvalue spectrum of a *linear* evolution operator, or transition matrix in the case of discrete states. The eigenvectors associated with the gaps in the spectrum indicate slow collective modes and provide natural targets for coarse-graining. In the hydrodynamic example of ∼ 10^23^ interacting molecules, these modes are the effective variables.

Here we seek such Markov dynamics from the time series of posture in the foraging behavior of the nematode worm *C. elegans*, an important model organism in genetics and neuroscience [8–1 For both the worm and animals generally, the collection of high-resolution behavioral data has been greatly accelerated by advancing techniques for pose estimation via machine vision [11–15], combined with computational and imaging improvements. Such measurement advances demand new behavioral understanding: analyses, models, and theory of posture-scale dynamics [4, 16, 17].

We implement a principled, generally-applicable framework which combines delay embedding with Markov modeling [6]. In this approach, we seek to overcome the partial observability of behavioral dynamics; variables which influence behavior but are instantaneously hidden become apparent over time and Markov predictability provides the quantitative measure of a self-determined system. Posture itself is a very complicated function of its underlying biological variables. In such situations, an initial expansion of dimensionality can simplify computations like function estimation and classification. We thus trade the complex modeling of a low-dimensional time series for the simpler modeling of a much higher-dimensional state space: the encoding of the unobserved degrees of freedom through time delays drastically simplifies our theory, leading to a powerful yet simple description of the emergent nonlinear Markovian dynamics.

While Markov approaches have an extensive history, perhaps most familiarly in Markov Chain Monte Carlo sampling of *equilibrium* distributions [18], substantially less attention has focused on a Markov encoding of actual *dynamics*, especially with a large number of states. Importantly, we note that there is no guarantee that our approach will work; for example, the number of necessary delays may be computationally prohibitive. But even this “failure” would provide important information about the memory of the system. On the other hand, if we are successful, we will be left with a finite set of observables that are approximately self-determined, measurable, and whose dynamics span the timescales that are relevant to the phenomena of interest. Such observables are likely to be biologically meaningful.

We find state variables for worm behavior that exhibit Markovian evolution across the multiple timescales of *C. elegans* foraging behavior: from fine-scale posture movements to “run-and-pirouette” strategies. Additionally, the macroscopic variables we reveal are not some baroque non-physical mathematical functions but rather correspond to interpretable behavioral motifs. We rediscover canonical behaviors from the rich history of *C. elegans* ethomics, as well as describe new ones. Each of these motifs is associated with its own characteristic timescale, and with them we provide a new hierarchical subdivision of behavior. We show how the dynamics of these macroscopic variables can be propagated through a model of the organisms physical interaction with the environment to accurately predict locomotion from posture. Finally, we dissect the function of these behavioral motifs by investigating their relation to the exploration and exploitation of food sources.

### SHORT-TIME BEHAVIORS AS MAXIMALLY-PREDICTIVE POSTURE SEQUENCES

On a 2D agar plate, worms move by making dorsoventral bends along their bodies [19]. At the shortest timescales (∼ 1 *s*) these traveling waves along the body give rise to forward, backward, and turning locomotion [20–22]. We show here that this organization emerges naturally from short posture sequences, formally a delay embedding [23–27].

We employ a previously analyzed dataset [28] composed of 35 minute recordings of 12 lab-strain N2 worms freely moving on an agar plate, sampled at *δt* = 1*/*16 s. From high-resolution videos, we measure the worm’s body posture using a rich low dimensional representation of the centerline, expressed as five “eigenworm” coefficients 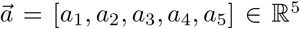, Fig. 1(a). We construct a maximally predictive sequence space [6] by stacking *K* delays of the posture time series, and increasing *K* until we have maximized the predictability of the resulting dynamics, as measured by the entropy rate, Fig. 1(b).

**FIG. 1.**
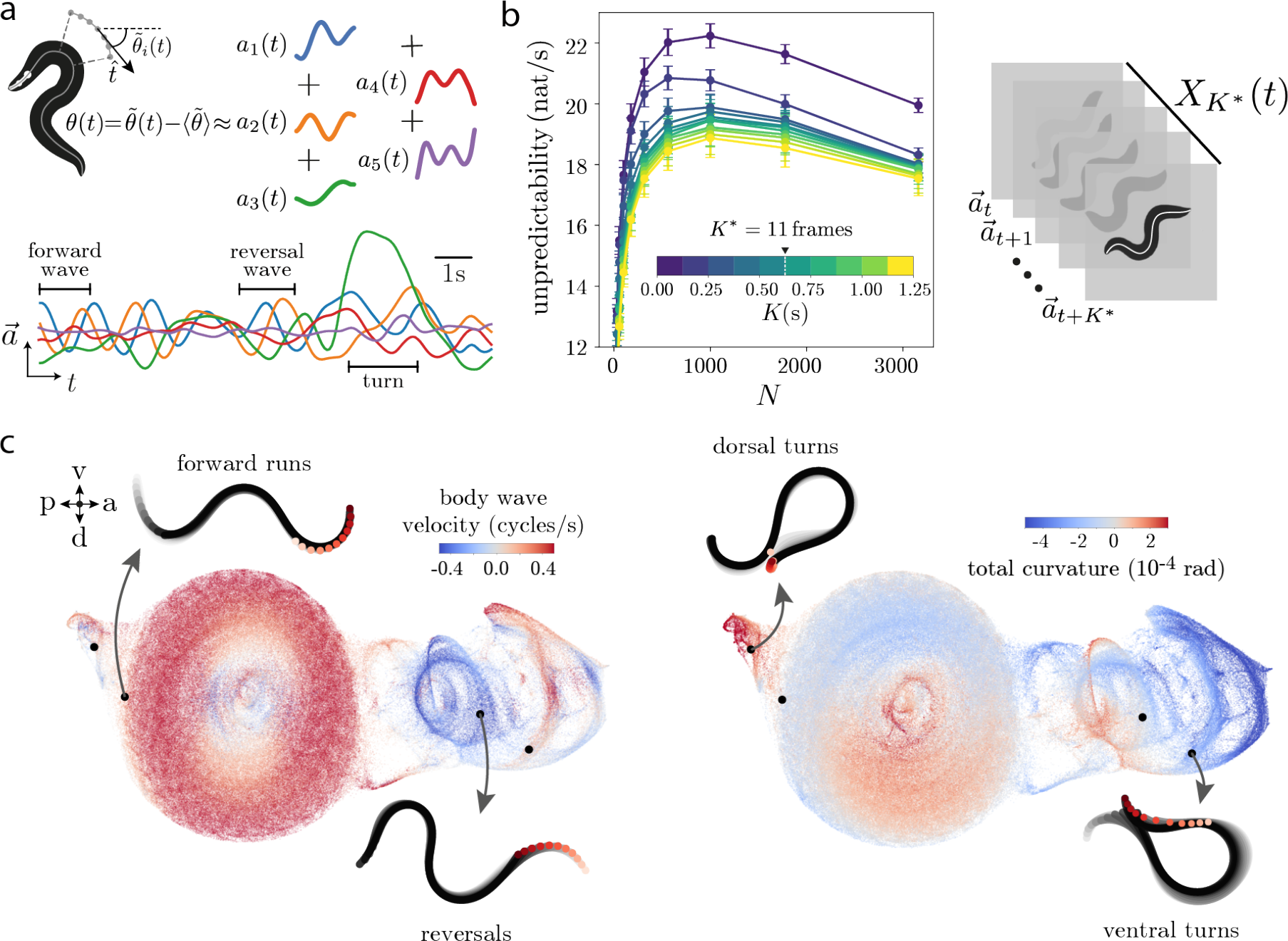
Maximally-predictive posture sequences reveal the space of short-time behaviors. (a-left) We represent the posture at each frame *t* using an “eigenworm” basis [20]. We extract the worm’s centerline and measure the tangent angles between body segments 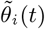. We then subtract the average angle to obtain a worm-centric representation 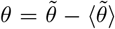, and project the mean-substracted angles onto a set of eigenworms [20], obtaining a 5D (eigenworm coefficient) time series 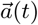. (a-bottom) A segment of the time series: in food-free conditions, the short time scale behavior roughly consists of forward and reversal waves (clearly visible as oscillations in *a*_1_ and *a*_2_), as well as sharp turns (characterized by large amplitude *a*_3_ [28]). (b) Entropy rate as a function of the sequence length *K* and number of partitions *N* . We partition each sequence space (indexed by length *K*) into N microstates using k-means clustering and compute the entropy rate of the resulting Markov chain. The curves collapse after *K* ∼ 0.5 s, indicating that the entropy rate is approximately constant meaning that there is no further gain in predictability by including more time delays. We choose *K*^∗^ = 11 frames = 0.6875 s to define a maximally predictive sequence space *X* _*K*∗_ . Error bars are bootstrapped standard deviations across worms. (c) We visualize *X*_*K*∗_ by projecting]onto two-dimensions using UMAP [29], and coloring each point by the body wave phase velocity 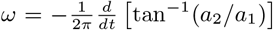 [20] (left) and the overall body curvature (right) obtained by summing the tangent angles *θ*_*i*_ along the body *γ* = Σ _*i*_ *θ*_*i*_. Each point on this space corresponds to a *K*^∗^ sequence of postures, and different short-time behaviors naturally correspond to different regions on the projection.

To estimate the entropy rate, we partition each sequence space (indexed by *K*) into *N* microstates using k-means clustering so that the worm’s posture dynamics now appear as transitions between microstates, a Markov chain. We approximate the entropy rate as that of the inferred Markov chain and choose the largest *N* before finite size sampling reduces the estimated entropy ^1^. This “maximum entropy” partitioning requires a large number of microstates but enables our model to be maximally expressive. After *K* ≈ 8 frames = 0.5 s and the entropy rate curves start to collapse, and we set *K*^∗^ = 11 frames = 0.6875 s to define the maximally-predictive sequence space *X*_*K* ∗_ ; lengthening the sequences does not increase predictability, Fig. S1(a). At *K*^∗^ we choose a partition of size *N* ^∗^ = 1000.

We visualize *X*_*K*∗_ by projecting into two dimensions using the UMAP manifold learning algorithm [29] (see Methods), Fig. 1(c). Color-coding according to the worm’s body wave phase velocity *ω* = 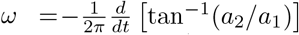, [20] Fig. 1(c, left), and overall curvature (obtained by summing the tangent angles along the body), *γ* = Σ _*i*_ *θ*_*i*_, reveals that distinct short-time behavioral motifs, corresponding to forward, reversal, and turning movements, naturally correspond to different regions of the maximally predictive state space, Fig. 1(c, right). In other words, while the instantaneous posture itself *-a*(*t*) is not enough to disentangle different behaviors, a point in the sequence space *X*_*K*∗_ (*t*) uniquely corresponds to a particular short-time behavioral motif.

### THE MARKOV WORM

To better understand behavioral dynamics in the posture sequence space 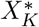 we trade individual trajectories for the evolution of the probability density, formally akin to the choice of a Langevin (trajectory) vs. a Fokker-Plank (ensemble) perspective [30, 31], and here we briefly sketch the mathematical framework. We expect the maximally predictive sequences to evolve according to 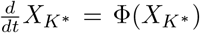, where Φ(*X*_*K*∗_) is a nonlinear noisy function. The corresponding evolution of the probability densities 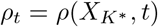 is 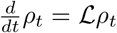, where *ℒ* is a differential operator that depends on the exact form of Φ. For a finite time step we have *ρ*_*t*+*τ*_ = *e* ^*ℒ τ*^ *ρ*_*t*_. Importantly, even when the trajectories evolve non-linearly, the probability dynamics are linear. We approximate this linear ensemble evolution *e* ^*ℒ τ*^ as a finite-dimensional Markov chain with *N* microstates *s*_*i*_ determined by partitioning the space of posture sequences, and a transition matrix *P* (*s*_*j*_(*t* + *τ*)|*s*_*i*_(*t*)) ≡ *P*_*ij*_(*τ*) ≈ *e* ^*ℒ τ*^ constructed by counting transitions between microstates *s*_*i*_(*t*) and *s*_*j*_(*t* + *τ*) after a delay *τ* [6],

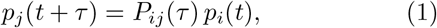

where *p*_*i*_(*t*) is the probability of observing state *s*_*i*_ at time *t* and we sum over repeated indices. Note that *P* is a stochastic matrix so that each transition probability *P*_*ij*_ ≥ 0 and Σ _*j*_ *P*_*ij*_ = 1 for all microstates *i*.

For the worm’s posture dynamics, as described in the previous section, we set the number of microstates as *N* = *N* ^∗^ = 1000 so as to maximize the amount of information with respect to the partitioning, Fig. 1(b). We set the transition time as *τ* ^∗^ = 0.75 s so that the relaxation times of the long-lived dynamics are approximately independent of *τ*, as rigorously true in a Markov process (see Fig. S1(b,c) and Methods). Despite the conceptual simplicity of our Markov chain model, Eq.1, we show that it accurately predicts *C. elegans* foraging behavior across scales, from fine-scale posture movements to long time scale transitions between behavioral states, Fig. 2.

**FIG. 2.**
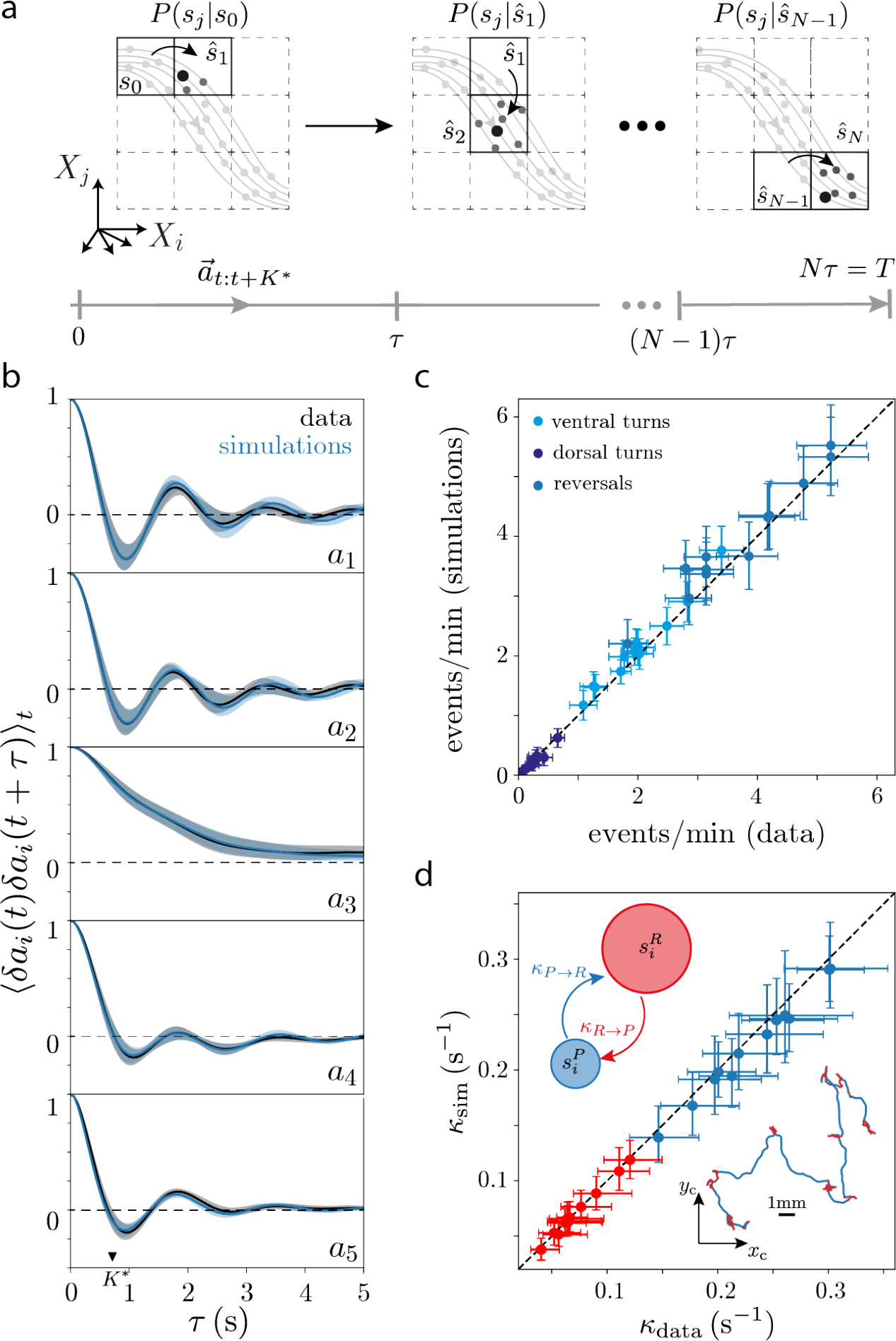
A Markov model captures posture dynamics across timescales. (a) Schematic of the simulation method. Starting from the initial microstate of each worm *s*_0_, the next microstate is obtained by sampling from *P*_0*j*_ (*τ* ^∗^). We add new microstates in the same fashion, resulting in a symbolic sequence of length *N* = *T δt/τ* ^∗^. Within each microstate, we randomly choose a point in the maximally-predictive sequence space, *X*_*K∗*_, and use this point to identify the associated sequence of body postures 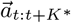. (b) The simulated posture dynamics accurately predicts the autocorrelation function of the “eigenworm” coefficient time series. Shaded regions correspond to 95% confidence intervals of the estimate of the mean autocorrelation function bootstrapped across worms. (c) The simulated posture dynamics accurately predicts the average rate of behavioral events across worms. We estimate the number of reversals, ventral turns and dorsal turns per unit time, and compare the result obtained from the data with simulations of each of the 12 recorded worms, finding excellent agreement. Error bars correspond to bootstrapped 95% confidence intervals on the estimates of the mean. (d) On longer time scales, worms transition between relatively straight “runs” and abrupt reorientations through “pirouettes” [33], which are combinations of reversals and turns (inset, bottom right). We assign each microstate to either a “run” or a “pirouette” by leveraging the inferred transition matrix to identify long-lived stereotyped sequences of posture movementc (see section **COARSE-GRAINING BEHAVIOR THROUGH ENSEMBLE DYNAMICS**). We then estimate the mean transition rates from the run to the pirouette state *κ*_*R*→*P*_ (red) and from the pirouette to the run state *κ*_*P* →*R*_ (blue) for each of the 12 recorded worms. Data error bars are 95% confidence intervals of the mean bootstrapped across run and pirouette segments, simulation error bars are 95% confidence intervals of the mean bootstrapped across 1000 simulations.

#### Predicting behavior across scales

Starting from the initial state of an individual worm, we simulate symbolic sequences by sampling from the conditional probability distribution *P*^*w*^(*s*_*j*_|*ŝ*_*i*_), Fig. 2(a), where *ŝ*_*i*_ is the current microstate, *s*_*j*_ are all possible future microstates after a time scale *τ* ^∗^ and *P*^*w*^(*s*_*j*_|*ŝ*_*i*_) is the *i*-th row of the transition matrix inferred for worm *w*. The result is a sequence of microstates with the same duration as the worm trajectories, but with a sampling time *δt* = *τ* ^∗^. From each microstate we can then obtain a nearly continuous time series of “eigenworm” coefficients 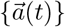 through the sequence of *K*^∗^ postures in each state *X*_*K*∗_ (note that *K*^∗^and *τ*^∗^ are quite close in this case). These dynamics are effectively diffusive in the space of posture sequences: hopping between microstates according to the Markov dynamics, followed by random selection from the set of posture sequences 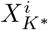 within each visited microstate *s*_*i*_. The posture time series generated through this procedure are nearly indistinguishable from the data, Fig. S2(a) and SI Movie 1. Quantitatively, the autocorrelation functions of the simulated time series, Fig. 2(b), capture the correlations observed in the data, and the distribution of mode coefficients agrees with the steady-state distribution, Fig. S3. In addition to the finescale posture dynamics, our model also predicts the rate at which forward movements are interrupted by biologically relevant behaviors [32] such as reversals, dorsal turns or ventral turns (identified by thresholding the body wave phase velocity [20] and the overall body curvature, see Methods), Fig. 2(c). Finally, at larger spatio-temporal scales the foraging random-walk can be coarsely split into forward “runs” interrupted by sharp “pirouettes”: sequences of reversals and turns used by the worm to reorient itself [33]. Here we identify “runs” and “pirouettes” directly from posture dynamics by using the inferred transition matrix to identify stereotyped sequences of states (see the following section COARSEGRAINING BEHAVIOR THROUGH ENSEMBLE DYNAMICS). As illustrated in the inset of Fig. 2(d), the identified states split the trajectory into “runs” and “pirouettes”. We estimate the kinetic transition rates from runs-to-pirouettes *κ*_*R*→*P*_ and from pirouettes-toruns *κ*_*P* →*R*_ and find close agreement between data and simulations across worms, Fig. 2(d).

#### Posture to Path

The accuracy of our Markov dynamics suggests the intriguing possibility that we may also recover the properties of foraging trajectories, i.e. the motility of the worm in 2D space. With such a bridge we could, for the first time, connect the neuromechanical control of posture with movement strategies such as optimal search. To do so, however, we must connect posture deformations with movement in the environment.

Following previous work [34], we approximate the interaction between the worm’s body and the viscous agar surface through resistive force theory (RFT) [35]. This phenomenological approach assumes that each segment along the body experiences independent drag forces. Despite its simplicity, this approximation has been successfully applied to predict the motility of various organisms in viscous fluids [36–38] and granular materials [39].

To propel the worm, we first reconstruct the skeleton positions in each frame **x**_*i*_(*t*) from the instantaneous tangent angles *θ*_*i*_(*t*) along each body segment *i*, Fig. 4(aleft). From these we derive the worm-centric velocities **v**_*i*_(*t*) = **x**_*i*_(*t* + 1) − **x**_*i*_(*t*) and displacements with respect to the center-of-mass position **x**_CM_, ∆**x**_*i*_(*t*) = **x**_*i*_(*t*) − **x**_CM_(*t*). This results in an expression for the underlying velocities at each body segment as a function of the measured worm-centric **v** and ∆**x** and unknown overall translational 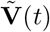 and angular 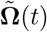 velocities. As in [34], we use linear resistive force theory to decompose the force acting on each body segment into tangent and normal components 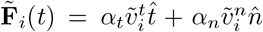, with drag coefficients *α*_*n*_ and *α*_*n*_, Fig. 4(a-middle). Using this approximation, we can then recover the unknown underlying velocities 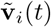 by imposing a zero net force and torque condition. The only free parameter in this model is the ratio between the normal and tangential drag coefficients *α* = *α*_*n*_*/α*_*t*_, which we infer by minimizing the distance between the reconstructed centroid trajectories and the real data (see Methods), Fig.S4(a). In agreement with the results of Keaveny et al. [34], we find that in such food-free conditions, the value of *α* that optimizes the reconstruction of centroid trajectories is *α*^∗^ = 30 (29, 31). Using such *α*^∗^, we then reconstruct the centroid path corresponding to the posture time series simulated according to our Markov chain, and show that these qualitatively resemble real worm trajectories, Fig. 3(a-right).

**FIG. 3.**
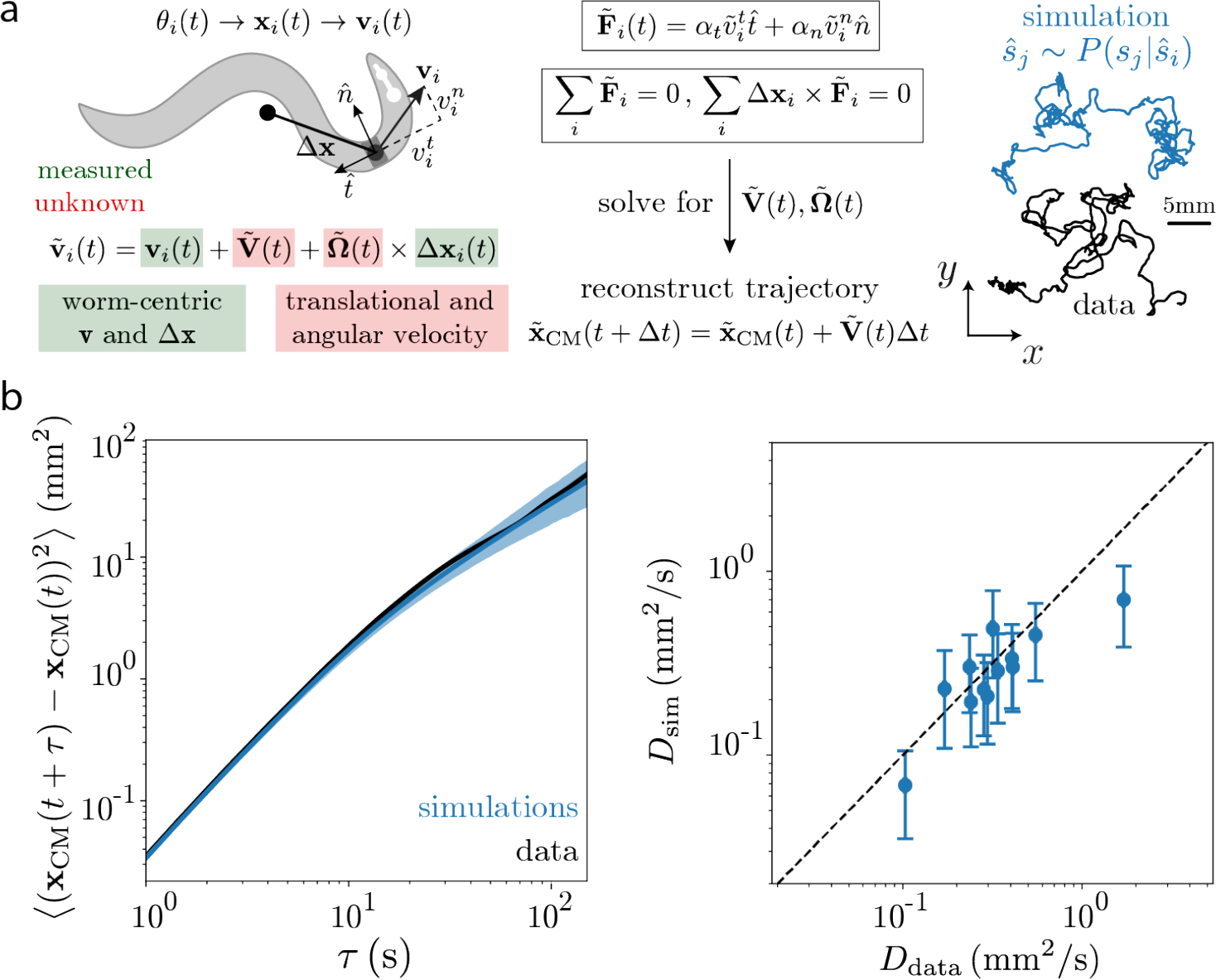
Recovering foraging trajectories from posture simulations. (a) We use resistive force theory (RFT) [34] to translate the simulated posture dynamics into movement. We simulate the Markov chain posture dynamics as described in Fig. 2. At each frame *t* of the simulated posture time series, we reconstruct the coordinates of the *i*-th segment of the skeleton, **x**_*i*_(*t*), from the tangent angles *θ*_*i*_(*t*), which are themselves a linear combination of eigenworms with the mode weights particular for each frame. The measured velocities, **v**_*i*_, in the frame-of-reference of the worm, correspond to subtracting the overall velocity 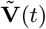 and angular velocity 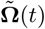 of the worm from the lab-rame velocities 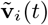, which are unknown. Following the results Ref. [34], we recover the lab-frame translational 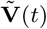 and angular 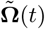 velocities by leveraging resistive force theory to equate the tangent and normal forces acting in each local segment to the local velocities and imposing a zero net-force and net-torque condition. The ratio between tangent *α*_*t*_ and normal *α*_*n*_ damping coefficients is the single free parameter *α* = *α*_*t*_*/α*_*n*_, which we find by minimizing the distance between simulated and real trajectories, Fig. S4(a). We show an example worm trajectory (black), as well as simulated trajectories reconstructed from posture time series generated from the operator dynamics (blue). (b-left) Mean square displacement of centroid trajectories obtained from model simulations (blue) and data (black) for an example worm. Simulation error bars are 95% confidence intervals of the mean bootstrapped across 1000 simulations. The results for all 12 worms analyzed here can be found in Fig. S4(b). (b-right) Effective diffusion coefficients obtained from simulations and from the data. We estimate *D* by fitting the slope of the mean square displacement curves in the range *τ* ∈ [60, 100] s, MSD(*t*) = 4*Dt*. Errors are standard deviations of the estimated diffusion coefficients across simulations.

**FIG. 4.**
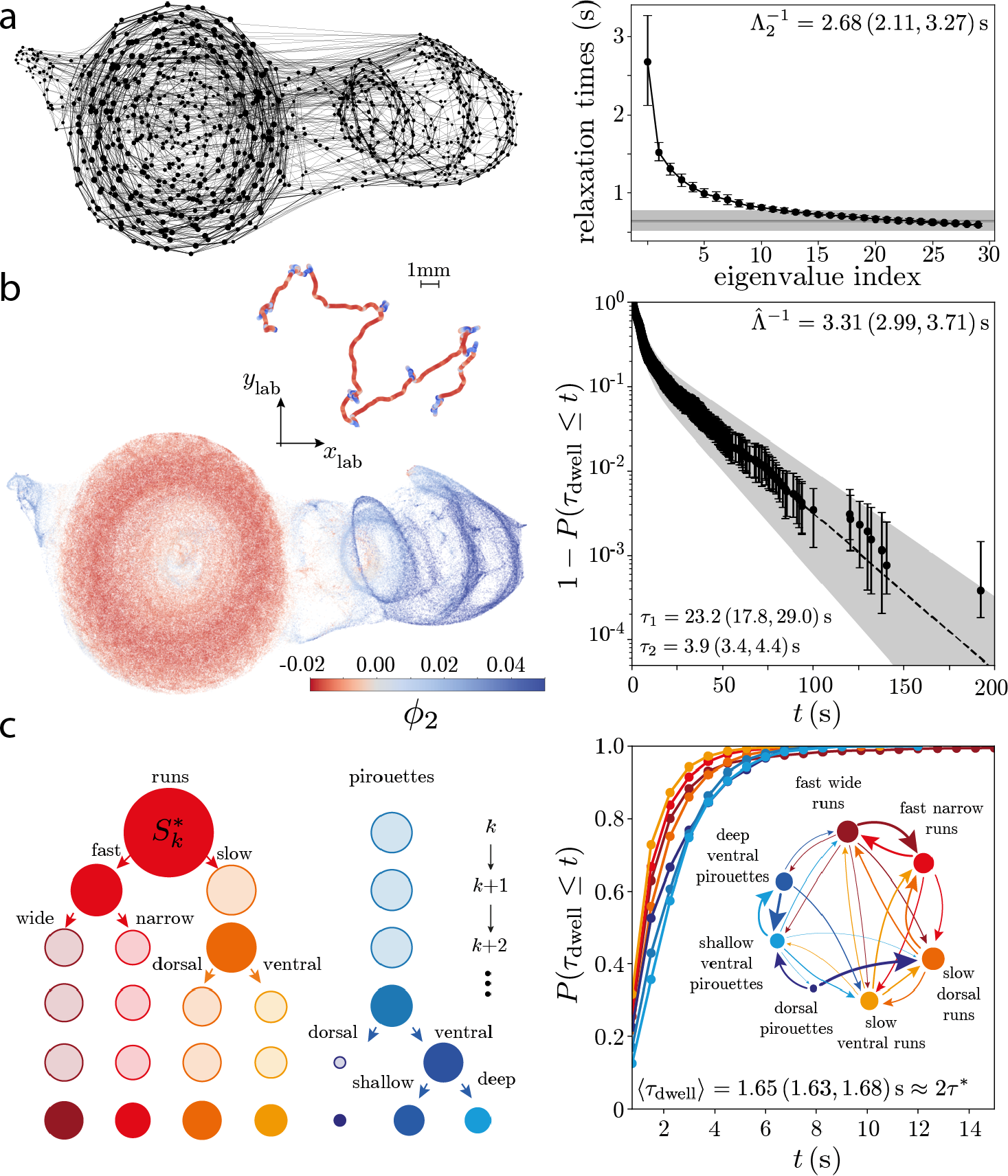
Slow modes for coarse-graining: from “runs-and-pirouettes” to stereotyped body waves. (a-left) Schematic of the inferred Markov chain. We partition the maximally predictive sequence space *X*_*K*∗_ (11 *×* 5 dimensions) into Voronoi cells, here represented as points in the 2-dimensional UMAP embedding space. The size of each point is proportional to the probability of visiting a given microstate *s*_*i*_, and linewidth corresponds to the probability of transitioning among distinct states after *τ* = 1 frame (we only show transitions with *P*_*ij*_ *>* 0.025 for simplicity). (a-right) Relaxation timescales obtained from the 30 eigenvalues with the largest real part of *P*_*ij*_ (*τ* ^∗^) with *τ* ^∗^ = 0.75 s. The horizontal gray bar is the largest non-trivial eigenvalue of the transition matrix computed from a shuffled symbolic sequence. Error bars are 95% confidence intervals bootstrapped across worms. (b-left) We color the sequence space and an exemplar centroid trajectory (inset) by the first nontrivial eigenvector of the reversibilized transition matrix *ϕ*_2_, which optimally separates metastable states [6, 49]. Positive values correspond to combinations of reversals and turns that reorient the worm, these are “pirouettes”, while negative values correspond to “forward runs”. (b-right) We use *ϕ*_2_ to identify two coarse-grained regions, which we denote as “macroscopic” behavioral states, and find that the complementary cumulative distribution function of the resulting dwell times 1 − *P* (*τ*_dwell_ ≤ *t*) is characterized by two time scales, which we extract by fitting a sum of exponential functions. The inferred time scales are in agreement with previous phenomenological observations of worm behavior [33], and result in a relaxation time 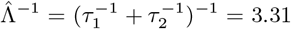 (2.99, 3.71) s [31] that agrees with the largest eigenvalue of 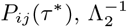, within error bars. The timescale errors and error bars are 95% confidence intervals bootstrapped across events. (c) The subdivision process. At each iteration step, we subdivide the metastable state with the largest measure, 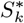, along the first nontrivial eigenvector obtained from the transition probability matrix conditioned only on the states within that metastable state [50]. This results in a top-down subdivision of behavior that follows the structure of an effective free energy landscape. The size of the circles represents the relative measure in each state. For interpretability, we stop at the 5th subdivision, yielding 7 “mesoscopic” states (characterized below). (c-right) Cumulative distribution of the mesoscopic state dwell times: the duration is short (≈ 2*τ* ^∗^). The inset shows a transition diagram in which the size of the nodes and the edge widths are proportional to the measure in each behavior and the transition probabilities, respectively (transition probabilities *<* 0.05 are not graphed for simplicity).

To further quantify the similarity between the centroid trajectories reconstructed from posture simulations and the data, we estimate the mean squared displacement MSD(*τ*) = ((**x**_CM_(*t* + *τ*) − **x**_CM_(*t*))^2^), which exhibits a transition between super-diffusive (nearly ballistic) and diffusive behavior between 10 s and 100 s [40–43], Fig. 3(b-left), Fig S4(b). The foraging trajectories corresponding to the operator-based simulations accurately capture the MSD across a wide range of scales, including the ballistic-to-diffusive transition. To further assess the quality of the simulations, we estimate an effective diffusion coefficient by fitting MSD = 4*Dτ* in the linear regime^2^ and find that, across worms, the resulting diffusion coefficients obtained from simulations closely follow the data, Fig. 3(b-right). Our results demonstrate that it is possible to go from microscopic posture dynamics to diffusive properties in a living organism.

### COARSE-GRAINING BEHAVIOR THROUGH ENSEMBLE DYNAMICS

As highlighted in Fig. 2, *C. elegans* foraging behavior exhibits multiple time scales: from the body waves that define short-time behaviors (e.g., forward, reversal, turns), to longer-time sequences (e.g., run, pirouette) that the worm uses to navigate its environment. Typically, these longer-time sequences have been identified phenomenologically by setting thresholds on heuristically defined quantities, as was done in Fig. 2(c). Here, we show that it is possible to reveal the multiple scales of *C. elegans* locomotion directly from the posture dynamics. Intuitively, stereotyped behaviors correspond to regions of the behavioral space that the animal visits often. In an analogy with statistical mechanics, we can imagine behavior as evolving on a complex potential landscape, where each well corresponds to a particular stereotyped behavior, and the barrier heights set the transition time scales. Such a picture emerges naturally when analyzing the dynamics through an ensemble approach, and we leverage our inferred Markov chain to directly identify metastable behaviors.

#### “Run-and-Pirouette”

The eigenvalues of the transition matrix provide direct access to the long time-scale properties of the dynamics, even when these are not directly apparent from the original trajectories or the equations of motion. The real part of the eigenvalues *{λ*_*i*_*}* of *P*_*ij*_(*τ* ^∗^) characterize the exponential relaxation to the steady state.

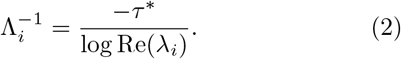

The spectrum of relaxation times is shown in Fig. 4(aright), and exhibits an isolated, longer-lived mode with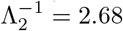. Although there are ≈ 10 significant modes, beyond the first mode, all others are indistinguishable from each other. To assess significance, we obtain a noise floor (horizontal line in Fig. 4(a-right)) by shuffling the symbolic sequence, reestimating the transition matrix, and computing its first nontrivial eigenvalue. In the limit of infinite data, there is only one surviving nonzero eigenvalue corresponding to the steadystate distribution (infinite relaxation time). The fact that the second largest eigenvalue is nonzero in the shuffle reveals finite-size effects that result in small deviations from the invariant density.

The eigenvectors corresponding to long-lived dynamics reveal reaction coordinates that capture transitions between “macroscopic” metastable sets [45, 46]. In Fig. 4(a-right), these are groups of microstates that transition more often within rather than between groups. The structure of these sets and the kinetics between them offer a principled coarse-graining, which is not imposed but rather follows directly from the ensemble dynamics. As in [6], we identify metastable sets through spectral analysis of the time-symmetric (reversibilized) transition matrix *P*_*r*_ (see Methods) [47, 48], whose second eigenvector *ϕ*_2_ provides an *optimal* subdivision of the state space into almost invariant sets [49].

To elucidate the meaning of the slow mode, we use *ϕ*_2_ to color-code the maximally-predictive sequence space, Fig. 1(c,d); Positive values (blue) generally align with negative phase velocities and large dorsal and ventral curvatures indicative of “pirouettes”, while negative values (red) correspond to positive phase velocities and low curvatures, indicative of “forward runs”. In the inset we show an example 10 minute long centroid trajectory color coded by the projection along *ϕ*_2_. Negative projections occur during “runs”, while positive values are found during abrupt reorientation events composed of sequences of reversals and turns. We thus obtain a slow reaction coordinate that captures the dynamics along a “run-and-pirouette” axis. The remaining eigenfunctions also reveal interpretable features of worm behavior, Fig. S5, albeit on a faster timescale.

To achieve a principled, data-driven, coarse-graining of the slow dynamics, we search along *ϕ*_2_ for a single threshold that maximizes the metastability of both resulting coarse-grained sets (see Methods) [6], Fig. S6, and we identify the resulting two macrostates as “run” and “pirouette”. In Fig. 4(b-right) we show that the complementary cumulative distribution of the resulting run lengths 1 − *P* (*t*_state_ *< t*) is roughly characterized by two time scales, Fig. 4(b-right), fit by a sum of exponential functions and in excellent agreement with previous phenomenological observations [33]. In addition, these transition timescales are related to the timescale of relaxation to the steady state distribution as 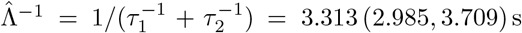 (2.985, 3.709) s [31], which agrees with the relaxation times of the transition matrix within statistical accuracy, Fig. 4(a-right). In our analysis “run-and-piroutte” kinetics emerge directly from worm-centric posture dynamics, without any positional information.

#### “Run(s)-and-Pirouette(s)”

While dividing the dynamics along *ϕ*_2_ identifies the longest lived states, splitting the effective free energy landscape along its highest energy barrier, what if there are important additional states within each metastable set? We use the transition matrix to perform a sequential subdivision of the posture embedding, revealing finer-scale states [50] (see Methods), Fig. 4(c). At each step, the metastable state with the largest measure is subdivided along the first nontrivial eigenvector of the reversibilized transition matrix conditioned solely on the microstates within the metastable state. This yields a subdivision of the posture embedding that obeys the structure of the free-energy landscape; at each iteration, we subdivide the system along the largest energy barrier within the highest measure basin. We note that our subdivisioning process proceeds *from the longest-lived states down* rather than from the shortest-lived states up, where the latter is more common in behavior coarse-graining approaches [51–53].

In the foraging behavior of *C. elegans*, beyond the initial division into runs and pirouettes (which we denote as “macroscopic” states), we further subdivide the dynamics into 7 “mesoscopic” interpretable states: 4 distinct run states and 3 subdivisions of the pirouette state Fig. 4(c),S The run state essentially splits into two fast states and two slower states, which can be distinguished either by the wave length of the body, or by having a particular bias towards the dorsal or ventral sides: the dorsally-biased slow state is akin to a head-casting state [54], while the ventrally-biased stated is akin to a “dwelling” state [55– with incoherent head and tail movements and no propagating wave [13]. On the other hand, the pirouette state neatly splits into dorsal turns, deep ventral *δ*-turns, and reversals followed by shallow Ω-turns.

These mesoscopic states that decorate the worm’s foraging behavioral landscape are short-lived, with a characteristic time scale of *⟨τ*_dwell_*⟩* = 1.65 (1.63, 1.68) s ≈ 2*τ* ^∗^, Fig. 4(c-right). The transition diagram between them, Fig. 4(c-right,inset), reveals the fine-scale organization of the worms’ foraging strategy. Further subdivisions result in even shorter-lived states, which are increasingly challenging to interpret.

### EXPLORING THE ROLE OF THE MESOSCOPIC STATES

In the data analyzed here, worms were grown in a food-rich environment, but then placed on food-free agar plates and allowed to move without restrictions. Under these conditions, the worm’s behavior has been qualitatively described as foraging [58, 59]. We apply our approach to better understand the role of the mesoscopic states in the worm’s search for food.

We use the Markov model to simulate *in silico* worms that are forced to remain in a particular mesoscopic state and the posture-to-path framework to investigate the properties of the trajectories resulting from the posture dynamics in each of the states. We can simulate trajectories that are much longer than those observed in the data, Fig. 4(c-right), allowing us to dissect how different states produce distinct large-scale tracks. In Fig. 5(a), we show simulated 10 min long trajectories for each of the 7 mesoscopic states. Notably, the difference in posture wavelengths exhibited by the two fast “run” states, Fig. S7(b), results in dramatically different trajectories, with the longer wavelength state (fast wide runs) resulting in overall straighter paths, and the shorter wavelength state (fast narrow runs) resulting in ventrally-biased curved trajectories with a diameter that is several times the body length and a period orders of magnitude longer than the body wave period, Fig. S8. Interestingly, the dorsally-biased slow state also results in loopy trajectories, but with a shorter diameter and faster recurrence time. In addition, the ventrally-biased “dwelling”-like slow state [55–57] with its frequent head retractions results in a denser sampling of a local patch. Finally, the three “pirouette” states result in a denser sampling of space and a reduced centroid displacement.

**FIG. 5.**
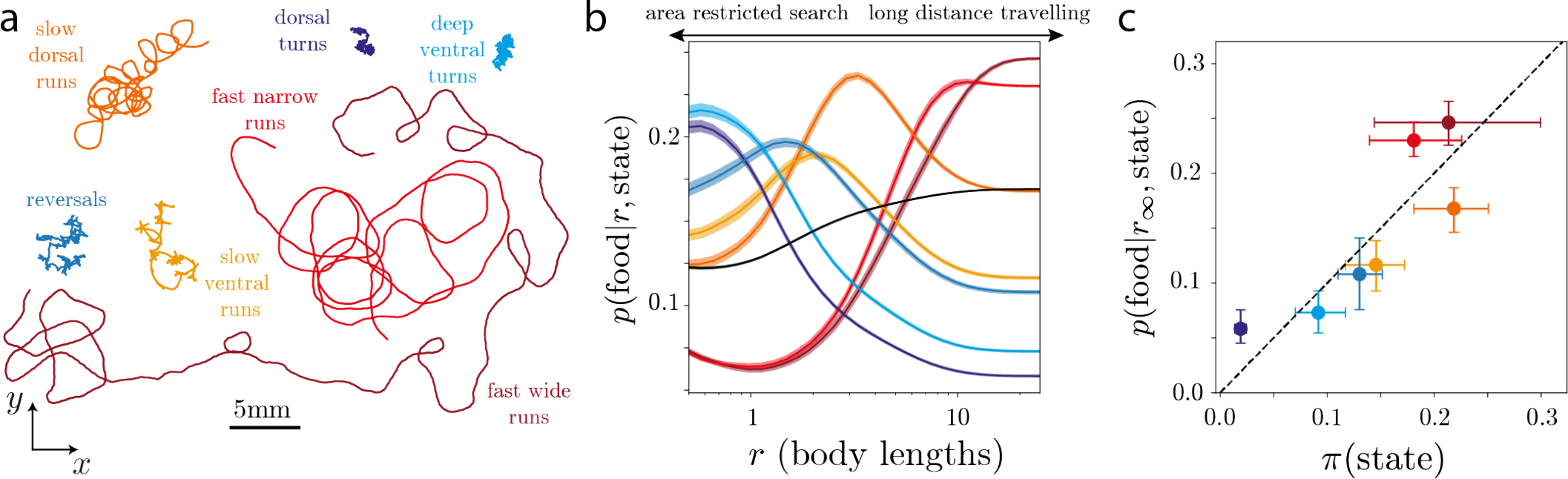
Leveraging our multiscale description to explore the role of behavioral states. We simulate long trajectories while forcing the worm’s dynamics to remain within a single mesoscopic state *S* obtained through a subdivion of the behavioral space into 7 states, Fig. 4(c). We proceed as in Figs. 2,3, but sample each new state according to a transition matrix built only within the partitions belonging to *S, ŝ*_*j*_ (*t* + *τ*) ∼ *P*_*S*_ (*s*_*j*_ |*ŝ*_*i*_(*t*)), *i, j* ∈ *S*. (a) Example 10 min long centroid trajectories of each state. (b) Probability of finding food in a uniform food patch of radius *r* for a particular state. We find that the most efficient behavioral states change depending on the radius of the food distribution: at short distances, “pirouette” and slow “run” states provide better chances of finding food, whereas at larger distances the two fast “run” states are most efficient. The black line represents the strategy of an average worm *p*(food|*r*, average worm) = Σ _*s*_ *π*(*S*)*p*(food|*r, S*), obtained as a weighted average over the probability of finding a worm in a particular mesoscopic state *S, π*(*S*). While the worm’s movement strategy is more likely to find food sources scattered at larger distances, the ensemble of mescoscopic states can be utilized for foraging success across scales. (c) The likelihood of finding food in a large uniform food patch for each of the states closely matches the probability of finding a worm in a particular state.

We next interrogate the efficiency of each of the 7 mesoscopic states at encountering food uniformly distributed within a disc of radius *r* around the initial position, Fig. 5(b), a simple but informative condition. We find that the “pirouette” states as well as the slow “run” states are most efficient at finding food at shorter distances, while at larger distances the two fast “run” states perform best. Such a differential use of behaviors is also seen in nature. Upon encountering food, *C. elegans*, as well as many other species, engage in area restricted search, which is characterized by shorter paths and a high frequency of large angle turns [42, 58, 60–65]. Conversely, upon removal from food, *C. elegans* lowers its turning rate [41, 66] to engage in global search or long distance travel [58, 62–65].

Remarkably, we find that instead of only using the most efficient behavioral state (“fast wide runs”), worms engage in a strategy that employs each mesoscopic state in a proportion that closely matches the relative efficiency of the different states at finding food uniformly distributed in a large patch (several body lengths), Fig. 5(c). This “probability matching” behavior has been observed across several species, including humans (see, e.g., [67–72]), and emerges naturally in “multi-armed bandit” situations in which agents must decide among different actions that yield variable amounts of reward without knowing *a priori* the relative reward of each action (see, e.g., [73]).

## DISCUSSION

We combine maximally-predictive short posture sequences with a Markov chain model to bridge disparate scales in the foraging dynamics of the nematode worm *C. elegans*. Rather than seeking low-dimensional descriptions of the data directly (e.g. [42, 57, 74–77]), we instead first *expand* in representation complexity: enlarging the variable of interest to include time in the form of posture *sequences* and constructing a maximum entropy partition to capture as much predictive information as possible. This expansion both in time and number of micostates is similar in spirit to that currently found in large language models, though our conceptual approach is dramatically simpler.

The maximally-predictive sequence space combines worm postures from roughly a quarter of the duration of a typical body wave, in agreement with previous work [22]. On longer timescales, the posture-based “run-and-pirouette” navigation strategy [33, 78] derived from the inferred Markov dynamics provide an accurate and principled coarse graining of foraging behavior, disentangling motions that are confounded by centroid-derived measurements (see e.g. [79]). This is particularly evident in our subdivision of the behavioral space. For example, we identify distinct “run” gaits that exhibit comparable centroid speeds, but are clearly distinguishable by the posture dynamics. Additionally, our top-down subdivision of behavior reflects the hierarchy of timescales in *C. elegans* foraging behavior [54]. Our approach systematically identifies such a control hierarchy from behavioral recordings alone, connecting posture timescales to “run-and-pirouette” kinetics. It will be interesting to investigate how the mesoscopic states identified here are controlled by the nervous system of the worm, and recent advances in experimental techniques that permit simultaneous neural and behavioral imaging in *C. elegans* provide an exciting path toward such discoveries [80–

The power of our modeling approach is in its simplicity; we bridge scales using a simple but effective Markov model, and this is only possible by recognizing and exploiting the mutual dependence between modeling and representation. Instead of directly modeling the posture time series (which can require higher-order and highly non-linear terms, see e.g. [20]), we search for maximally predictive states such that a simpler Markovian description can nevertheless accurately predict behavior. These emergent Markov dynamics offer a promising and powerful demonstration of quantitative connections across the hierarchy of movement behavior generally exhibited by all organisms [76, 85].

By finely partitioning the space of posture sequences, we encode continuous nonlinear dynamics through a Markov chain with a large number of states. This is analogous to building a hidden Markov model (HMM), but one in which the “hidden” states are actually observable (through time delays of our observations), and for which there is a one-to-one correspondence between “hidden” states and emitted symbols: each observation in the posture sequence *-a*(*t*) uniquely determines the state *X*_*K*∗_ (*t*). While HMMs are commonly used in behavioral analysis (see e.g. [13, 86]), they are rarely built with so many states and with the goal of correctly predicting dynamics. In particular, most approaches employ a small number of discrete behavioral states, where the number of states is a hyperparameter of the model and the discretization is not unique. In contrast, we let the data *reveal* the “hidden” states through time delays, and set the discretization so as to maximize predictive information. In this sense, the HMM we build is unique: by revealing the *hidden* dynamics through time delays, the “hidden” states are uniquely determined by the observations, making the HMM unifilar [87]. In other words, the “hidden” states themselves have a very definite meaning in our approach: we effectively group together “pasts” that have equal predictability over the future up to an *E*-resolution (set by the number of partitions 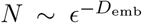, where *D*_emb_ is the intrinsic embedding dimension of the dynamics), approximating the system’s *causal states* [88]. This set of states together with the resulting Markov chain effectively constitutes an *E*-machine [89], the minimal maximally-predictive machine. Any other HMM in which hidden states are not *causal* returns models that severely overestimate the complexity of the dynamics. In addition, even though we start with a large transition matrix, we can coarse-grain it by identifying which states commonly follow each other in time to generate stereotyped sequences. In this way, instead of imposing discrete states from the start (as is common with HMM approaches), we first identify a large number of predictive causal states and only then leverage the resulting Markovian dynamics to identify coarse-grained stereotyped behaviors.

Our information theoretic framework also frees us from the constraint of linearity that is commonly imposed in graphical models applied to animal behavior (such as autoregressive hidden Markov models [90]). In particular, while the stereotyped states found through such models are encoded by linear dynamics, the states we identify can exhibit much more complex nonlinear dynamics, allowing us to capture longer time-scale structures in behavior.

Are Markov models enough to capture the richness of animal behavior more universally? It is important to distinguish between two sources of non-Markovianity. The first one is general and is simply induced by the fact that time series data are typically only a partial observation of the full dynamical state [6, 7]. Projecting the full unobserved dynamics onto a subset of observable degrees of freedom inherently results in non-Markovian dynamics for the measured variables [2, 3, 91, 92]. Such an “under-embedding” might result in apparent memory when naively constructing behavioral states. The second source of non-Markovianity, which is less trivial and likely ubiquitous in behavior, derives from the fact that there may be “hidden” latent variables that modulate behavior over timescales comparable to the measurement time. In this case, the steady-state distribution itself is changing slowly over time, rendering the dynamics explicitly non-stationary [5, 93].

A relevant demonstration of non-stationarity is provided by the adaptive changes in pirouette rate seen in the behavior of *C. elegans* upon removal from a food-rich environment [41, 42, 58, 62]. This adaptation is present in the data we analyze and is not captured by our Markov model, Fig. S9. To characterize such non-ergodic latent variables requires explicitly time-dependent Markov models, which we leave for future work. We note, however, that our coarse graining can be easily extended to capture non-stationary dynamics through the discovery of *τ* -dependent coherent sets that identify *moving* regions of the state space that remain coherent within a time scale *τ* [94–99].

Particularly interesting future directions include the analysis of even longer dynamics in *C. elegans* [13, 100– 102], where we expect to be able to extract longer-lived behavior strategies, such as the minutes-long transitions between “roaming” and “dwelling” states in food-rich environments [55–57]. Our modeling approach can also be used as means to obtain a deeper understanding of the effects of genetic, neural, or environmental perturbations on the multiple scales of *C. elegans* behavior. Indeed, the inferred transition matrices are a powerful phenotype that encapsulates multiple scales of *C. elegans* foraging behavior, and has the power to reveal how behavior is affected by a given perturbation. Of particular relevance for the study of long timescales in behavior would be to focus on mutations that impair neuromodulatory pathways and are thus likely to impact the spectrum of relaxation times of the inferred Markov chain.

The effectiveness of our Markov model at capturing the nonlinear dynamics of *C. elegans* body pose, combined with the ability to translate those spatiotemporal dynamics into movement, have allowed us to investigate how different behavioral states result in distinct ways of exploring the environment at much larger scales. Our analysis recovered the two main foraging modes exhibited by *C. elegans*: one that combines different pirouette states and slow runs resulting in a local search, and another one that mostly leverages fast run states to search for food more globally [42, 58, 62, 63].

We also discovered that the relative use of different behavioral states closely follows the relative efficiency of each state in food discovery. In fact, instead of using the behavioral state that would maximize its chances of finding food in its environment, worms match their strategy with the relative efficiency of each state. Interestingly, such a strategy, termed probability matching [71], or Thompson sampling [103], is a well-studied heuristic solution for the multi-armed bandit problem, a game in which different actions have variable rewards that are *a priori* unknown to the player, whose goal is to maximize total pay-out. Evidence for probability matching in decision making tasks has been previously demonstrated in experiments in animals and humans [104, 105], and is an active area of research in cognitive science of decision making [106].

While this strategy seems “irrational” in the context of maximizing reward in a fixed environment, that is not the condition in which worms have evolved: in ecologically-relevant situations the environment changes over time, rendering the distribution of rewards nonstationary and the subsequent sampling events correlated. Interestingly, reinforcement learning agents that have been evolved in a changing environment also develop probability matching strategies [107]. In addition, it has been shown that optimal Bayesian learners engage in probability matching when they expect sampling events to have temporal dependencies [108], as is the case in most ecologically relevant scenarios in which samples of the environment are not independent, but exhibit temporal correlations (at least as a result of the actions taken by the agent). This suggests that probability matching may reflect an exploration-exploitation tradeoff that robustly maximizes reward in an ever-changing environment. Our results indicate that *C. elegans* may implement such a heuristic in its foraging strategy. If the worm is indeed probability matching, it may have a way of storing its estimate of the current probability of success for each strategy (which may be reflected the dynamics itself). It will be fascinating to look for signatures of this in situations where we can experimentally adjust the pay-out probabilities. It may also be possible in a simple organism like *C. elegans* to estimate metabolic costs to utilize each behavioral state [109]. The multitude of genetic tools [110], the ability to image neurons in behaving animals [80–84] and to quantify behavior using our methods may make *C. elegans* an ideal system to look inside of an organism performing the dynamic loop of experiencing the world, making decisions based on those observations and its internal model of the world, and updating that internal model based on the outcomes of those decisions and their effect on the environment.

## METHODS

### Software and data availability

Code for reproducing our results is publicly available: https://github.com/AntonioCCosta/markov_worm/. Data can be found in [111].

#### *C. elegans* foraging dataset

We used a previouslyanalyzed dataset [20], in which N2-strain *C. elegans* were imaged at *f* = 32 Hz with a video tracking microscope on a food-free plate and downsampled to *f* = 16 Hz to incorporate coiled postures [28]. Worms were grown at 20°*C* under standard conditions [112]. Before imaging, worms were removed from bacteria-strewn agar plates using a platinum worm pick, and rinsed from *E. coli* by letting them swim for 1 min in NGM buffer. They were then transferred to an assay plate (9 cm Petri dish) that contained a copper ring (5.1 cm inner diameter) pressed into the agar surface, preventing the worm from reaching the side of the plate. Recording started approximately 5 min after the transfer and lasted for 2100 s, for a total of *T* = 33600 frames. Each frame is converted into a 5dimensional “eigenworm” representation 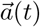 by projecting the local tangent angles along the worm’s centerline onto an “eigenworm” basis [20], Fig. 1(a).

### Maximally predictive states

Given the measurement time series, *-*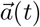, with *t* ∈ *{δt*, …, *Tδt}* and 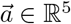, we build a trajectory matrix by stacking *K* time-shifted copies of 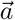, yielding a (*T* − *K*) *× Kd* matrix *X*_*K*_. For each *K*, we partition the candidate state space and estimate the entropy rate of the associated Markov chain (see below). We choose *K*^∗^ such that ∂_*K*_*h*(*K*^∗^) ∼ 0, which defined 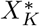 as the maximally predictive states [6], Fig. S1(a).

### State space partitioning

We partition the state space into *N* Voronoi cells, *s*_*i*_, *i* ∈ *{*1, …, *N }*, through kmeans clustering with a k-means++ initialization using scikit-learn [113].

### Transition matrix estimation

We build a finite dimensional approximation of the Perron-Frobenius operator using an Ulam-Galerkin discretization [46]. In practice, given T observations, a set of *N* partitions, and a transition time *τ*, we compute

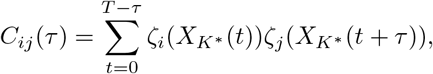

where ζ_*i*_(*x*) are the Ulam basis functions, which are characteristic functions

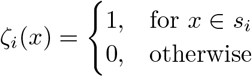

set by the k-means clustering. The maximum likelihood estimator of the transition matrix is obtained by simply row normalizing the count matrix,

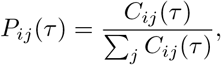

which yields an approximation of the Perron-Frobenius operator.

### Invariant density estimation

Given a transition matrix *P*, the invariant density is obtained through the left eigenvector of the non-degenerate eigenvalue 1 of *P, πP* = *π*: *π*_*i*_ is the probability of finding the system in a partition *s*_*i*_.

### Short-time entropy rate estimation

Given a number of partitions *N* and a sampling time scale *τ* = *δt*, we estimate the Markov transition matrix *P* and the corresponding invariant density *π* as detailed above and compute the short-time entropy rate as,

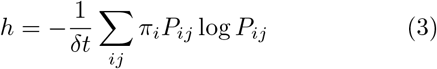

### Two-dimensional UMAP embedding

We use the UMAP embedding [29] as a tool to visualize the maximally predictive states of *C. elegans* posture dynamics. In a nutshell, the UMAP algorithm searches for a low dimensional representation of the data that preserves its topological structure. We use a publicly available implementation of the algorithm found in https://github.com/lmcinnes/umap, within which we chose the Chebyshev distance metric to compute distances in the high-dimensional space, `n neighbors`=50 nearest neighbors and `min dist` =0.05 as the minimum distance.

### Matrix diagonalization

The high dimensionality and the sparsity of the transition matrices for large *N* results in numerical errors when using a naive estimator for the full spectrum of eigenvalues. In addition, since we are interested in the longest lived dynamics, we focus on finding only the *n*_modes_ largest magnitude real eigenvalues using the ARPACK [114] algorithm.

### Choice of transition time *τ* ^∗^

We choose *τ* ^∗^ such that the resulting Markovian dynamics approximate the long-term behavior of the system accurately, as in [6]. In practice, we find the shortest transition time scale after which the inferred implied relaxation times reach a plateau, Fig. S1(b,c). For *τ* too short, the approximation of the operator yields a transition matrix that is nearly identity (due to the finite size of the partitions and too short transition time), which results in degenerate eigenvalues close to *λ* ∼ 1: an artifact of the discretization and not reflective of the underlying dynamics. For *τ* too large, the transition probabilities become indistinguishable from noisy estimates of invariant density, which results in a single surviving eigenvalue *λ*_1_ = 1 while the remaining eigenvalues converge to a noise floor resulting from a finite sampling of the invariant density. Between such regimes, we find a region with the largest time scale separation which also corresponds to the regime for which the longest relaxation times, Eq. (2), are robust to the choice of *τ*, Fig.S1(b,c). For further discussion see [6].

### *C. elegans* posture simulations

At each iteration, we sample from the conditional distribution given by the inferred Markov chain *P* (*s*_*j*_(*t* + *τ* ^∗^)|*s*_*i*_(*t*)) to generate a symbolic sequence sampled on a timescale *τ* ^∗^. We then randomly sample a state space point *X* _*K*∗_ within the partition *s*_*i*_, and unfold it to obtain a sequence of postures 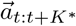 at each *τ* . We can thus generate artificial posture time series with the same duration as the experimental time series (35 minutes), but with a missing frame every *τ* ^∗^ frames (the gap between *K*^∗^ and *τ* ^∗^), which we interpolate across using a cubic spline with scipy’s `interpolate` package [115], and smooth with a cubic polynomial and a window size of 11 frames using the `signal.savgol filter` package from Scipy [115]. We then take the simulated *-a*(*t*) time series and transform it back to the tangent angles at each body segment *θ*_*i*_(*t*) using the “eigenworms” [20].

### Estimating the rate of reversals, dorsal and ventral turn events

Reversal events where identified as segments in which the absolute value of the worms’ overall curvature *γ*(*t*) = Σ_*i*_ *θ*_*i*_(*t*) was |*γ*| *<* − 3 *×* 10^−4^ rad and the body wave phase velocity 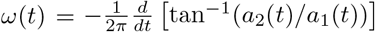 [20] was *ω <* −0.2 cycles s^−1^ for at least 0.5 s. Ventral and dorsal turns were identified as segments where the overall body curvature was either *γ <* −3.5 *×* 10^−4^ rad or *γ >* 3.5 *×* 10^−4^ rad, respectively, for at least 0.5 s.

### Resistive force theory simulations

We recover the rigid body motion from the tangent angle time series using linear resistive force theory, as in [34]. We approximate the forces acting independently on each body segment as

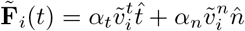

where 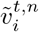 are the tangent and normal components of the velocity at each segment *i*, which can be written in terms of the velocity and displacements measured after subtracting the overall rigid body motion,

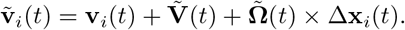

Then, by imposing a zero net-force and net-torque condition at each frame,

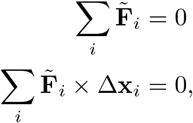

we obtain a system of linear equations that for a given *α* = *α*_*n*_*/α*_*t*_ can be solved for the components of the worm’s velocity 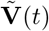 and angular velocity 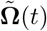 [34]. From these we can integrate the path taken by the worm’s body to obtain a reconstructed 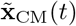.

We optimize the single free parameter *α* by comparing the reconstructed trajectories with the real worm trajectories 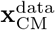, Fig. S4(a). In particular, we minimize the maximum distance between 100 s trajectories randomly sampled from the dataset 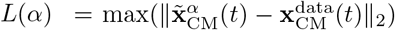, *t* ∈ [*t*_0_, *t*_0_ + 100 s]. To minimize *L*(*α*) we use the Nelder-Mead algorithm through the `scipy.optimize` library of Scipy [115]. The software to translate posture into path can be found in https://github.com/AntonioCCosta/markov_worm/, and follows closely the implementation of [34].

### Metastable states

Metastable states correspond to collections of short-time movements that typically follow each other in time to give rise to stereotyped sequences. Leveraging our previous work [6], we search for metastable states along the slowest mode of the reversibilized dynamics [47]. As shown in [49], the second eigenvector *ϕ*_2_ of a time-reversibilized transition matrix *P*_*r*_ provides an *optimal* subdivision of the state space into almost invariant sets. In practice, we estimate *P*_*r*_ as

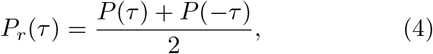

where,

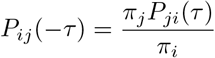

is the stochastic matrix governing the time-reversal of the Markov chain. The first non-trivial (*λ <* 1) right eigenvector of *P*_*r*_, *ϕ*_2_, allows us to define macrostates as collections of microstates *s*_*i*_,

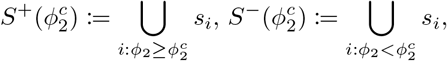

where 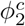 is a threshold that is chosen to maximize the metastability of a set. We measure the metastability of each set *S* by estimating how much of the probability density remains in *S* after a time scale *τ*,

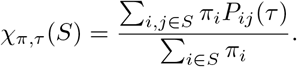

To estimate the overall measure of metastability across both sets *S*^+^ and *S*^−^, we define

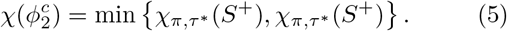

which we maximize with respect to 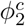. Metastable states are then defined with respect to the sign of 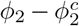. See [6] for further details and applications to known dynamical systems. In Fig. S6 we show the overall coherence measure as a function of *ϕ*_2_ for the worm data.

### Operator-based state space subdivision

We leverage the notion of relatively coherent sets [50] to subdivide the state space. However, instead of subdividing both metastable state at each iteration *k*, we identify the state with the most measure 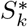 and build a new transition matrix only with partitions belonging to that state,

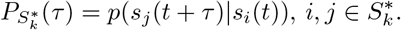

From 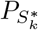 we proceed as before: we compute the stationary distribution of 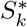 through the first left eigen-vector of 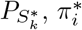, build the corresponding reversibilized transition matrix 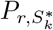 and identify relatively metastable states through its first non-trivial eigenvector by maximizing Eq. (5) where *π*_*i*_ and *P*_*ij*_(*τ*) are replaced by their relative counterparts 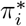 and 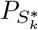 .

### Simulating posture-to-path within mesoscopic behavioral states

To generate a centroid trajectory within a given state, we construct a transition matrix among the partitions corresponding to each of the mesoscopic states identified in Fig. 4(c). We then proceed as in Figs. 2,3 to generate both posture time series and centroid trajectories. We first generate a symbolic sequence by sampling states according to the corresponding transition probability matrix *ŝ*_*j*_(*t*+*τ*) ∼ *P*_*S*_(*s*_*j*_|*ŝ*_*i*_(*t*)), *i, j* ∈ *S*. From the symbolic sequence, we then sample a time series segment 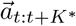 within each sampled partition, and use resistive force theory to translate the resulting *θ*(*t*) time series into locomotion. In this way, we can simulate posture and centroid trajectories for *in silico* worms that are forced to remain within a particular mesoscopic behavioral state for an arbitrary amount of time.

### Probability of finding food as a function of distance and behavioral state

We estimate the likelihood of finding food in a given radius *r* by estimating the fraction of the area within a disc of radius *r* covered by the worm’s body during 100 s trajectories, taking the worm’s width to be 5% of its length. We then normalize these area fractions by the total across states, obtaining the *p*(food|*r*, state) showed in Fig. 5(b).

## ACKNOWLEDGEMENTS

We thank Massimo Vergassola and Federica Ferretti for comments. This work was supported by OIST Graduate University (TA, GJS), a program grant from the Netherlands Organization for Scientific Research (AC, GJS), by the Herchel Smith Fund (DJ), and by Vrije Universiteit Amsterdam (AC, GJS). GJS acknowledges useful (in-person!) discussions at the Aspen Center for Physics, which is supported by National Science Foundation Grant PHY-1607611.

## SUPPLEMENTARY MATERIAL

**FIG. S1.**
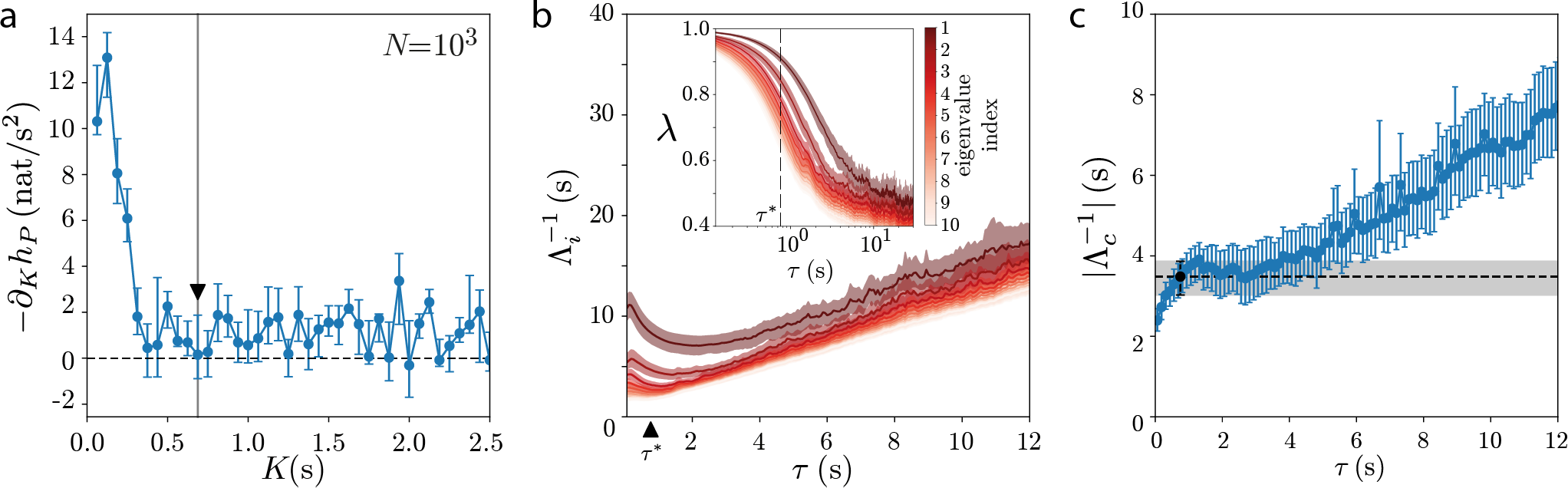
Details of the Markovian approximation of the *C. elegans* posture dynamics through maximally predictive states. (a) - Change in short-time entropy rate as a function of delays *K* for *N* = 1000 partitions. The entropy rates reaches a plateau after *K* 2: 0.5 s and we choose *K*^∗^ = 11 frames = 0.6875 s. Error bars represent 95% confidence intervals bootstrapped across worms. (b) - The ten largest relaxation timescales of the reversibilized transition matrix as a function of transition time *τ*, and corresponding eigenvalues (inset). For *τ* → *δt* the transition matrix is nearly the identity matrix (within *τ* most transitions occur within each partition), resulting in nearly degenerate eigenvalues close to 1 and an overestimation of the relaxation timescales of the reversibilized dynamics. On the other hand, when *τ ≳* 2: 5 s the dynamics is mostly mixed, meaning that the transition matrix is composed of near copies of the steady-state distribution. In this regime, the eigenvalues of *P*_*r*_, *λ*_*i*_, become approximately constant, and therefore 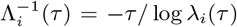 grows linearly with *τ* . Between these two regimes, the relaxation timescales are approximately constant, and this robustness to *τ* is indicative of Markovian dynamics. We choose *τ* ^∗^ = 0.75 s as the shortest *τ* ^∗^ consistent with Markovian dynamics. Error bars are 95% confidence intervals bootstrapped across worms. (c) - The reversibilized transition matrix provides an optimal partition into almost invariant sets (see Section **COARSE-GRAINING BEHAVIOR THROUGH ENSEMBLE DYNAMICS** for details), but the resulting kinetics does not necessarily capture the underlying dynamics. In fact, the obtained relaxation times are only an upper bound to the true relaxation timescales of the locally irreversible dynamics. To directly probe the Markovianity of the underlying slow dynamics, we estimate the relaxation times for the non-reversibilized coarse-grained transition matrix, which should not change with *τ* when the dynamics is Markovian. We approximate the slow relaxation dynamics by using the metastable states to build a two-state, coarse-grained Markov chain *P*_*c*_, which necessarily has only real eigenvalues *λ*_*c*_ ∈ R. The corresponding relaxation time is then obtained through 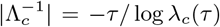. In general, we find that the regime in which 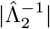 from *P*_*r*_ is constant (b) overlaps with regime in which 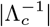 is also constant. In addition, while 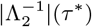 from *P*_*r*_ overestimates the expected |Λ^−1^| from “run” and “pirouette’ transition rates, Fig. 4(b), the timescales obtained from 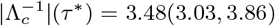 are comparable to the ones estimated from the entire Markov chain in Fig. 2(a) and accurately predict the hopping dynamics. Error bars are 95% confidence intervals bootstrapped across worms.

**FIG. S2.**
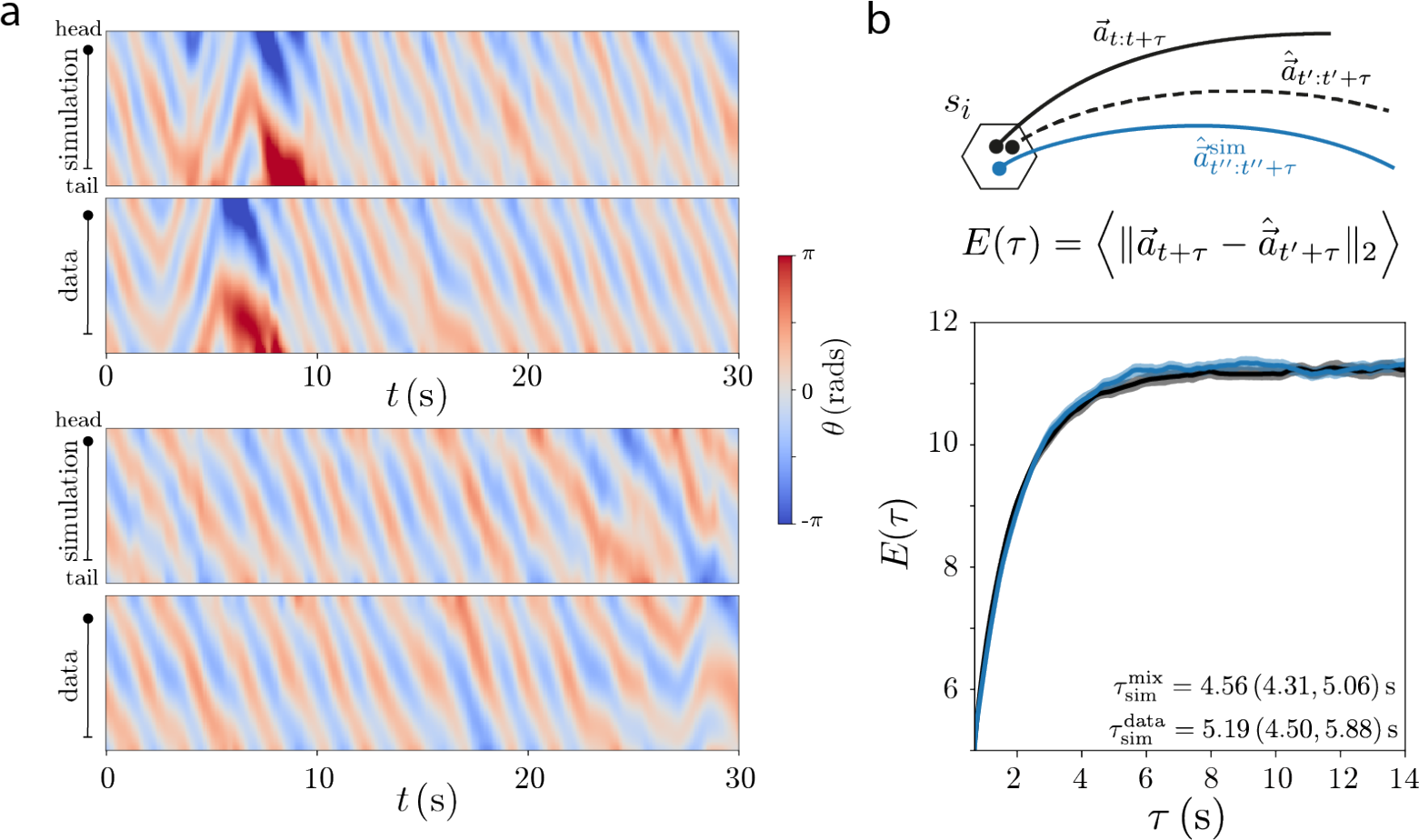
Details of the Markov chain simulations. (a) - Two illustrative curvature vs. time plots comparing simulations with data. As expected due to the unpredictable nature of the dynamics [22], the quality of the predictions worsens as time progresses. Nonetheless, the structure of the dynamics is well preserved, making it hard to know *a priori* which of the two time traces is the data and which one is a simulation. (b) - We assess the predictive power of the Markov model by estimating how prediction errors grow over time. We define *E*(*τ*) by estimating the average distance between two trajectories starting within the same partition: *t* is chosen at random and *t*^*’*^ *≠ t* chosen such that 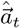*i* belongs to the same partition as 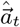; the expectation value is then taken over multiple samples of *t*. We compare *E*(*τ*) estimated from simulations (blue) against sampling a trajectory from the data starting from the same partition (black). As summary statistics, we compute the time it takes before predictions completely mix, obtaining 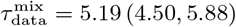 for the data, just slightly higher than 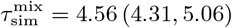 from simulations. In practice, we estimate the average distance between two randomly sampled points 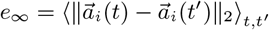, and find the time it takes for *E*(*τ*) ≤ 0.95*e*_∞_.

**FIG. S3.**
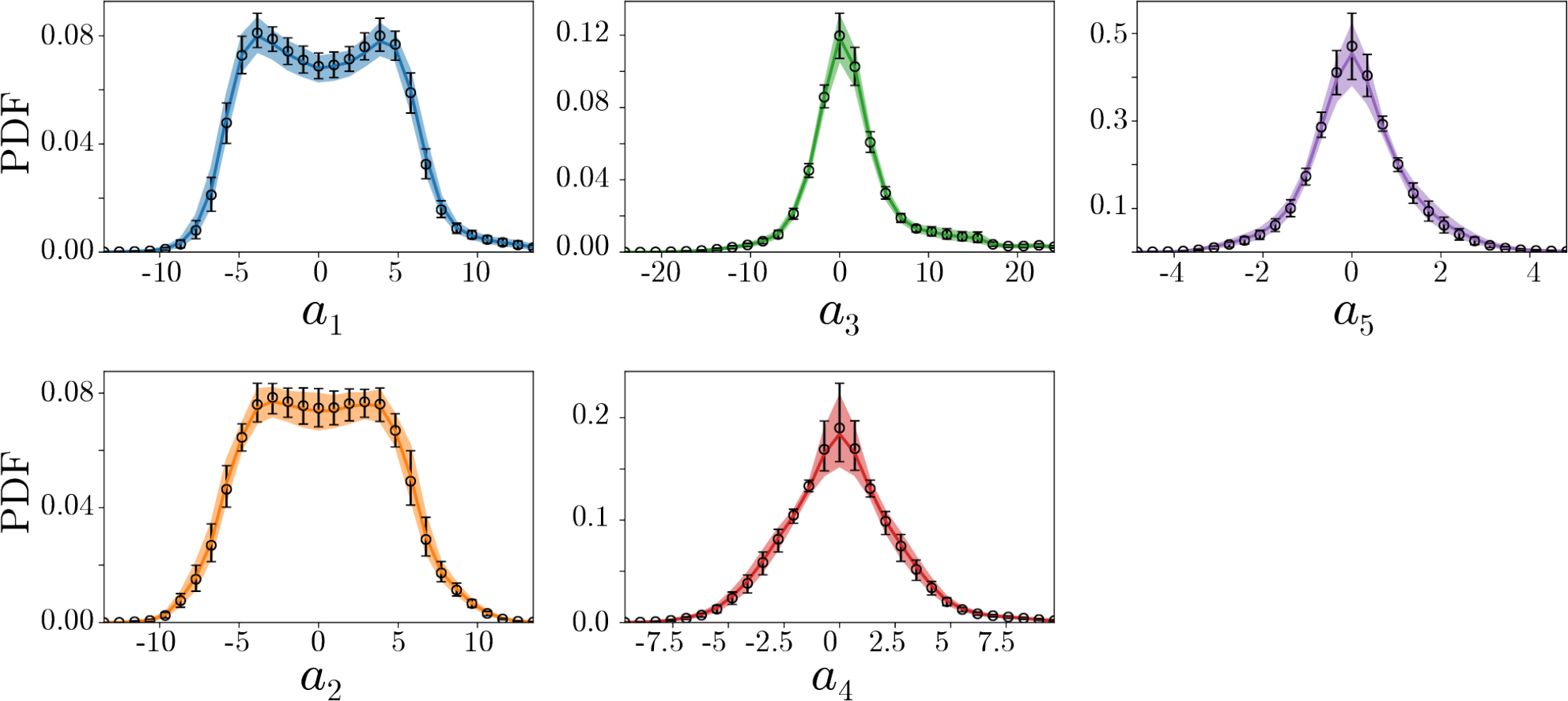
Probability Density Function (PDF) of the “eigenworm” coefficients show a tight agreement between the data (colors) and simulations (black error bars), indicating that the inferred dynamics capture the steady-state distribution. Error bars and shaded areas correspond to 95% confidence intervals bootstrapped across worms.

**FIG. S4.**
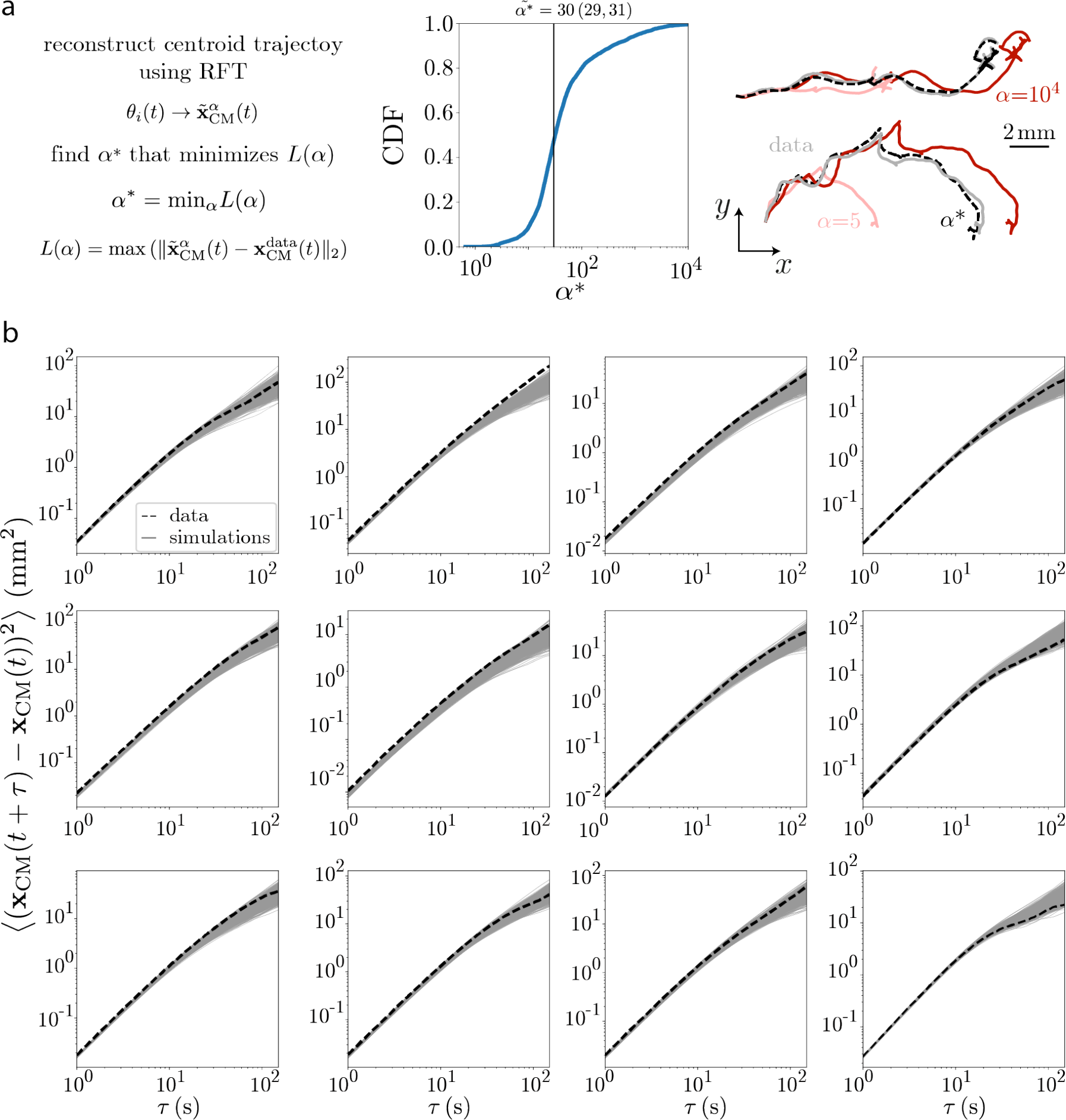
Details of the posture-to-path simulations. (a) Optimizing the free parameter *α* = *α*_*n*_*/α*_*t*_ in RFT (see Methods for details). We define a loss function *L*(*α*) as the maximum distance between the real worm trajectory and a reconstructed one, and sample randomly from 100 s segments. We the find the optimal *α*^∗^ through the Nelder-Mead algorithm (see Methods). The resulting distribution of *α*^∗^ is broad, as shown through the cumulative distribution function (CDF), with a median of 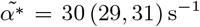. On the right, we display two example trajectory reconstructions for different values of *α*. For *α* = 5 (comparable to experimental measures of [116]), RFT typically results in a substantial undershoot of the observed trajectories (as observed in [34]). For the no-slip condition, *α* » 1, the resulting trajectories overshoot the real worm trajectories, indicating that some degree of slip is needed to accurately predict worm trajectories. With values of *α*^∗^ ≈ 30 we get an accurate reconstruction of the worm trajectories. (b) Mean square displacements for the data (dashed line) of each worm, as well as for 1000 centroid trajectory simulations generated from symbolic sequences simulated with the Markov model (gray). By fitting a linear function in the interval *τ* ∈ [60, 100] s we obtained the effective diffusivity estimate of Fig. 4(b-right).

**FIG. S5.**
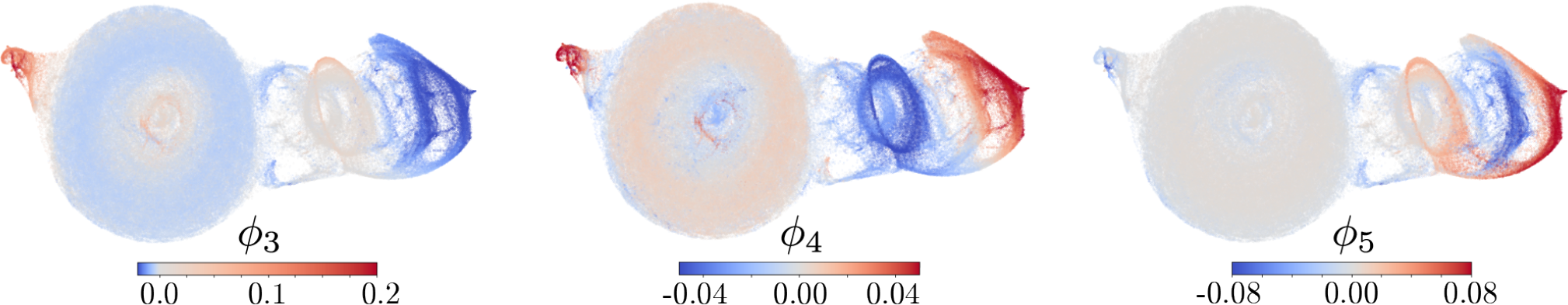
Example eigenvectors of the inferred transition matrix of *C. elegans* posture dynamics. While we focus primarily on *ϕ*_2_, there is also important information in the remaining long-lived eigenfunctions. To illustrate them, we color code the maximally predictive state space by the projection along the 3 following eigenvectors, *{ϕ*_3_, *ϕ*_4_, *ϕ*_5_*}*, which are organized according to their relaxation times 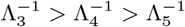. *ϕ*_3_ mostly captures an overall bias towards performing dorsal or ventral turns, *ϕ*_4_ decouples shallow pirouettes from recurring deep turns, and *ϕ*_5_ separates shallow turns following a pause from reversals that are followed by deep *δ*-turns. Notably, these eigenfunctions highlight stereotyped behavioral sequences on different timescales.

**FIG. S6.**
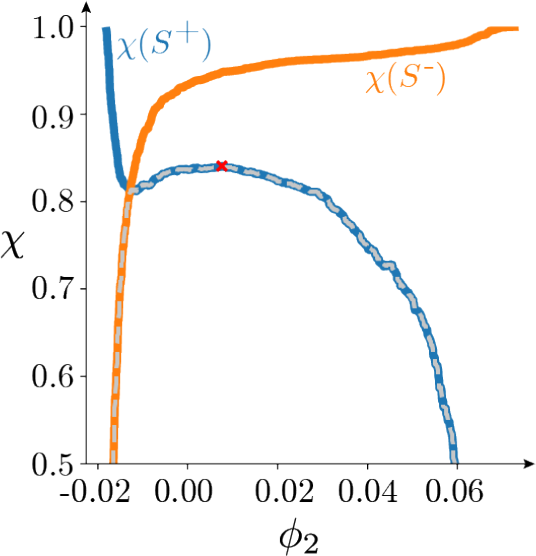
Coherence measure used to define the metastable states. We define metastable states by maximizing the overall coherence, Eq. (5), of the two macroscopic states obtained by partitioning the state-space along *ϕ*_2_ (see Methods for details). We here plot the coherence of each set (orange and blue), as well as the overall minima across sets *χ*, Eq. (5) (gray dashed line). The maximum of *χ* is highlighted with a red cross and indicates the value of *ϕ*_2_ that defines the metastable states, 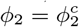.

**FIG. S7.**
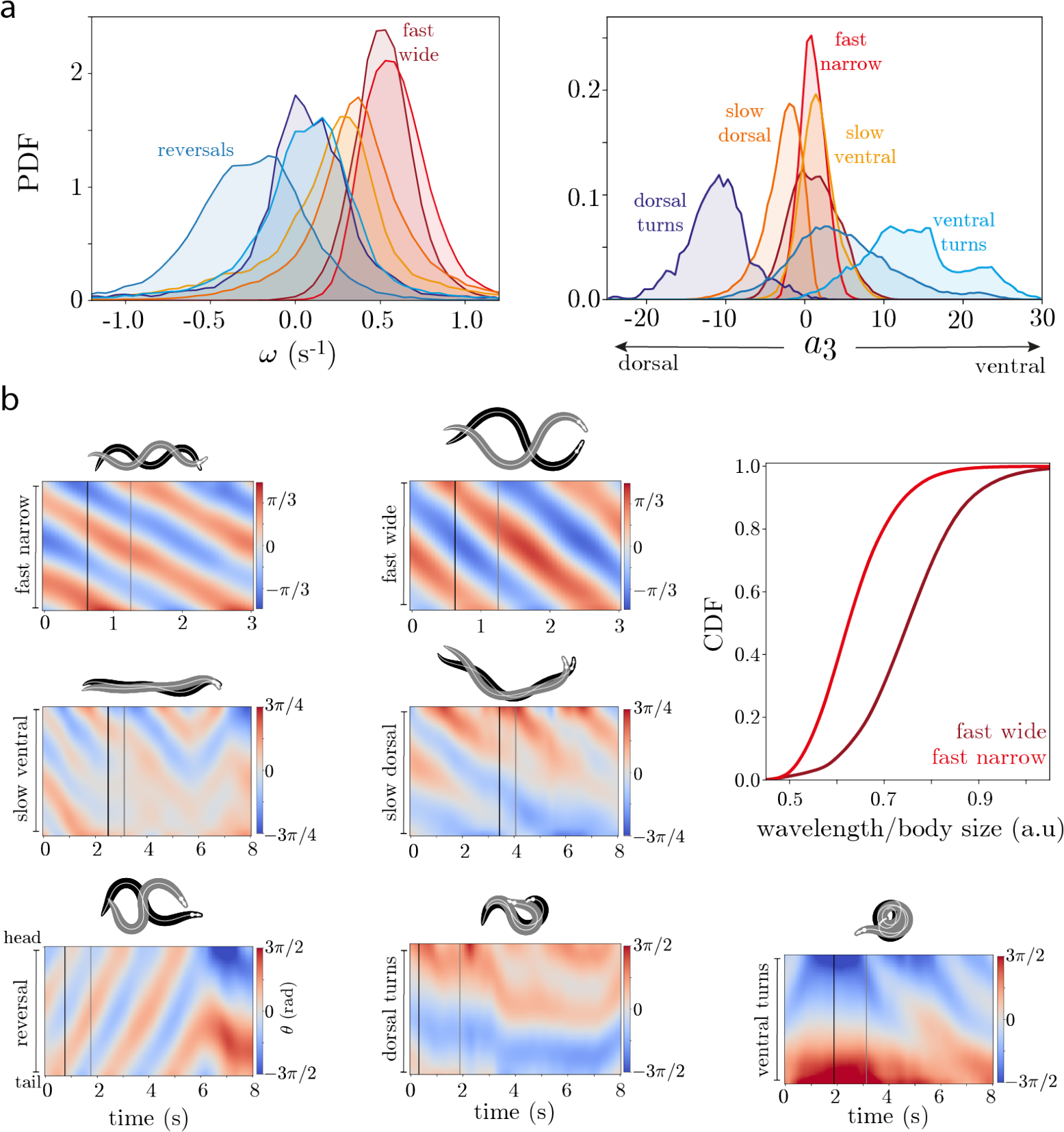
Characterization of the 7 mesoscopic states obtained through a top-down subdivision of the *C. elegans* (**posture sta**)te space. (a) - Probability Distribution Function (PDF) of the body wave phase velocity 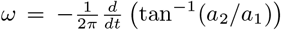 (left) and the dorso-ventral turning amplitude (right), as measured through the third “eigen- worm” coefficient *a*_3_ [20, 28]. The state labels were given based on these probability distributions. (b) Example kymographs of the local tangent angles *θ*_*i*_ as a function of time in each of the states, as well as example postures sampled at different time points (black and gray vertical lines). The two fast states (top) correspond to distinct gaits with different wavelengths (right), while the slow states (middle) lack a coherent body wave traveling from head to tail. Instead, the dorsally-biased slow states exhibits short timescale head-casting behavior [54], while the ventrally-biased state exhibits incoherent body motion with partial reversal and forward waves akin to a dwelling state [55].

**FIG. S8.**
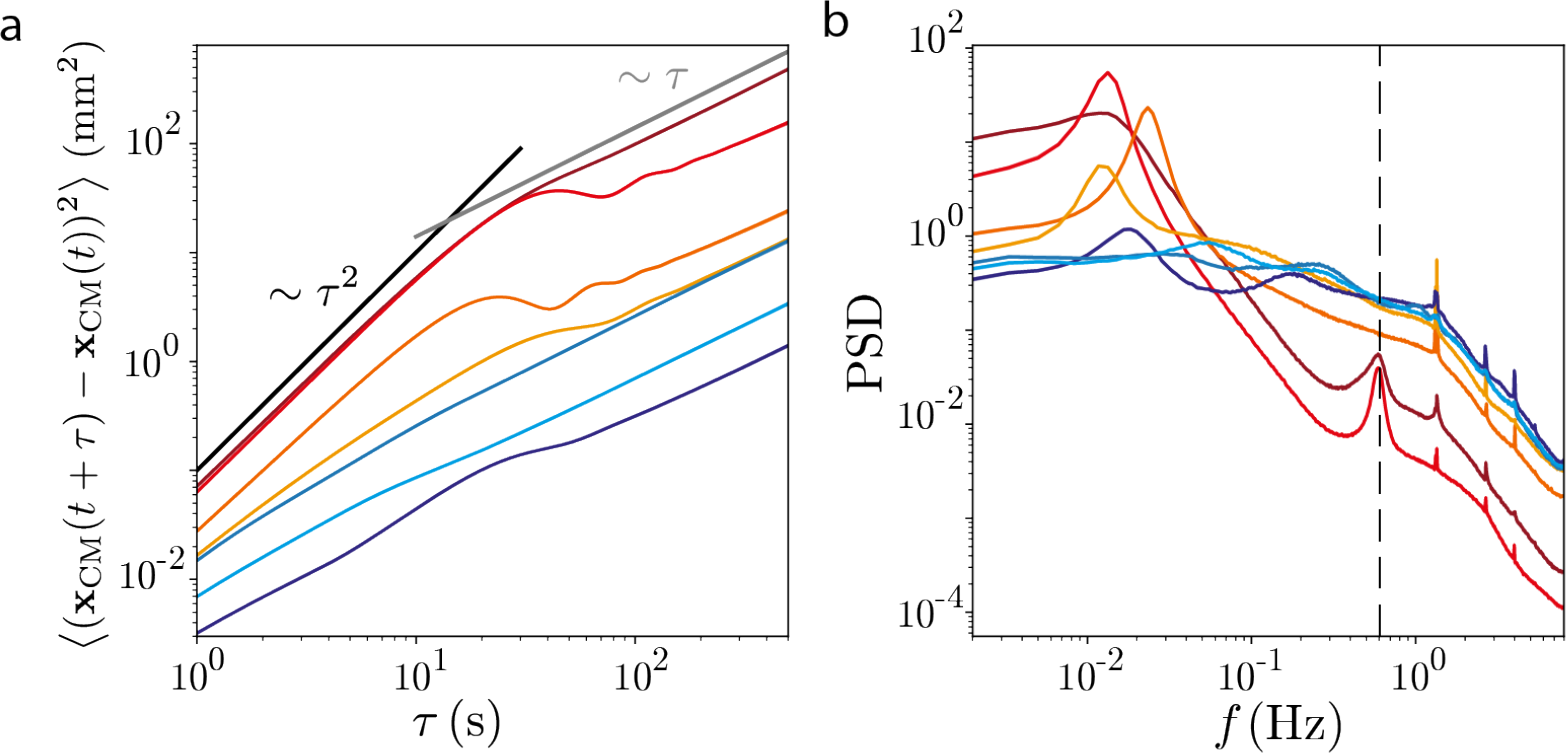
Statistical properties of the trajectories simulated for each of the 7 mesoscopic states obtained through a top-down subdivision of the *C. elegans* posture state space. (a) - Mean square displacement (MSD) of the center-of-mass trajectories for each of the 7 states revealed in Fig. 4(c). Trajectories in the different states exhibit clearly distinct statistical properties: while “run” states generally exhibit a transition between super-diffusive behavior (MSD ∼ *τ*^*β*^, *β >* 1) at short times and diffusive (or sub-diffusive) behavior (MSD ∼ *τ*^*β*^, *β* ≤ 1) at large times, the “pirouette”’ states are mostly diffusive even at short times. In addition, some of the states exhibit non-trivial fluctuations in the MSD that result from the quasi-periodic loops observed in the trajectories shown in Fig. 5(a). (b) - Power spectral density of the velocity bearing angle *η*, 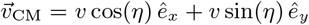, where 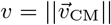. Besides the fast oscillations due to the body wave dynamics (the vertical dashed line represents the body wave phase velocity in the fast wide run state), some states also exhibit low frequency peaks due to the loopy nature of the trajectories, which recur on a time scale orders of magnitude longer than the body wave period. The power spectral density was estimated using Welch’s method [117] implemented through the signal.welch package from Scipy [115] with a Hann window and 10 min long trajectory segments. The error bars corresponding to 95% confidence bootstrapped across 1000 simulations for each state are too small to show.

**FIG. S9.**
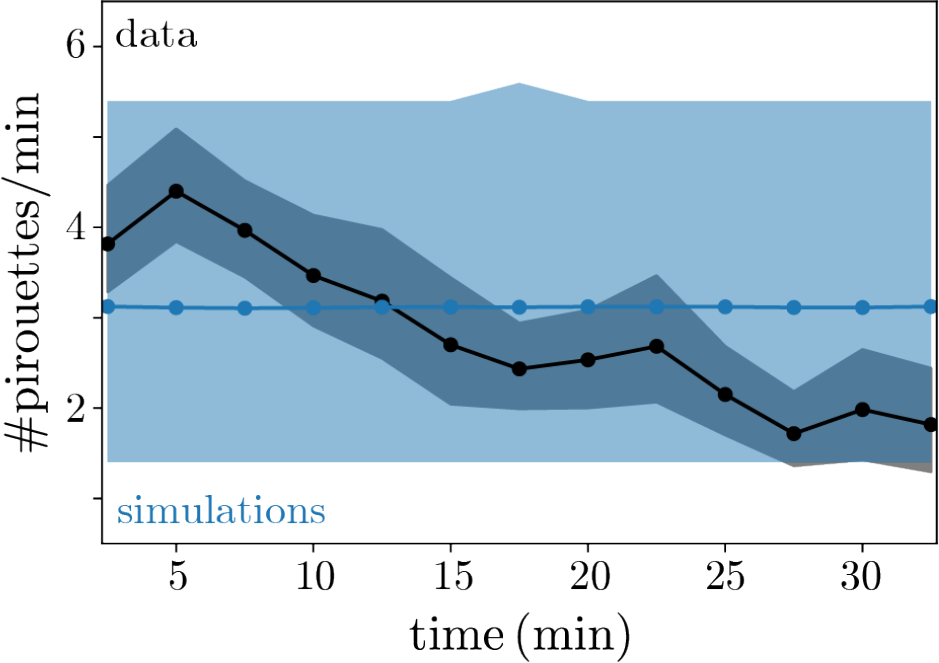
The rate of pirouettes changes with time on the food-free plate, a non-stationary dynamics not captured in our Markov model. We estimate the rate of pirouettes as a function of time from the data (black), and find that it slowly decreases, reflecting a change in search strategy possibly as a result of updates to the animal’s prior over the food distribution. Our simulations (blue) do not capture this slow change in the pirouette rate, as that would require an explicit time dependence in the transition probability matrix. Error bars correspond to 95% confidence bootstrapped across the 12 worms in our dataset. To estimate the rate of pirouettes, we coarse-grain the dynamics into “runs” and “pirouettes” as in Fig. 4, and find the number of pirouette events per minute in sliding 5 min windows with 2.5 min overlap. Since the sampling time of the Markov dynamics is *τ* ^∗^, we discard pirouette events with a duration shorter than *τ* ^∗^ from this analysis.

SUPPLEMENTARY MOVIE 1 (click to download). **The simulated posture dynamics is virtually indistinguishable from the real worm data**. We simulate a worm using the Markov model procedure of Fig. 2, and compare its posture dynamics with data starting from the same microstate. We show the curvature over time (left) for real (bottom) and simulated (top) worms, which are *a priori* indistinguishable from each other. We also show the corresponding RFT reconstructed skeletons (right), after subtracting the centroid position.

SUPPLEMENTARY MOVIE 2 (click to download). **Illustration of the posture to path simulations**. We simulate a worm using the Markov model procedure of Fig. 2, and translate its posture dynamics into movement using the resistive force theory approach of Fig. 3.

## Footnotes

1 The entropy rate of the Markov chain should not be confused with the Kolmogorov-Sinai (KS) entropy, which is an intrinsic property of the dynamics. Indeed, the KS entropy can also be estimated using our approach, yielding accurate estimates that agree with the sum of positive Lyapunov exponents [6, 22].

2 We note that on longer time scales the MSD exhibits the behavior of a confined random walk due to the rigid boundaries of the agar plate, which makes it non-trivial to accurately estimate the diffusion coefficient [44]. We fit the diffusion coefficient in the regime *τ* ∈ [60, 100], which corresponds to a time scale within which finite-size effects are negligible and the mean squared displacement is approximately a linear function of *τ*, MSD ∼ *τ*.

